# Pancreatic α-cells are functionally heterogeneous and sex-dependently regulated by neighboring endocrine cells

**DOI:** 10.1101/2025.08.23.671946

**Authors:** Ya-Chi Huang, Marjan Slak Rupnik, Guy A. Rutter

## Abstract

Glucose and paracrine regulation on α-cells, particularly with respect to sex differences, remains unclear. Hence, we imaged islets of GluCre:GCaMP6f^fl/fl^ mice in pancreatic slices, additionally loaded with a red Ca²⁺ dye, to precisely interrogate Ca²⁺ dynamics in α-cells and the adjacent β- and δ-cells. During a glucose ramp (1.8–10.8mM), α-cell Ca²⁺ oscillations were heterogeneous, antiphasic to β-cells, and inversely correlated with δ-cells. Selective inhibition of insulin receptor (InsR) and somatostatin receptor subtypes-2 and 3 (SSTR2/3) shifted the majority of α-cells to hyperactivity, delayed the decline of α-cell activity and onset of synchronized β-cell Ca²⁺ oscillations during the ramp. While female α-cells exhibited greater sensitivity to SSTR2/3 inhibition, male α-cells were more responsive to InsR blockade. Under complete SSTR2/3 and InsR antagonism, α-cells exhibited elevated Ca²⁺ oscillations but remained glucose-dependent. We conclude that intra-islet coordination fine-tunes α-cell glucose responses, with sex-specific paracrine signaling potentially shaping glucagon profiles in metabolic disease.

## Introduction

Glucagon, secreted by pancreatic α-cells, is a key hormone that counteracts the actions of insulin to prevent hypoglycemia. Under physiological conditions, glucagon secretion is potentiated at low glucose (<5mM) and suppressed at high glucose (>5-7 mM)^1^. The tight regulation of α-cell glucagon secretion, together with the coordinated control of insulin and somatostatin release from the neighboring β- and δ-cells respectively, is essential for maintaining normoglycemia. In diabetes, however, the regulation of α-cell activity is impaired^2^. Thus, in type 2 diabetes (T2D),

α-cells hyper-secrete glucagon, leading to excessive hepatic glucose production and worsening hyperglycemia. Conversely, in type 1 diabetes (T1D), α-cell glucose sensing mechanisms are compromised. Patients with T1D consequently experience postprandial hyperglycemia and face the life-threatening risk of hypoglycemia during periods of fasting.

Despite its critical role in glucose homeostasis, the regulation of α-cell glucagon secretion is incompletely understood. It is, for example, unresolved whether α-cells sense glucose directly^1,3–6^, or indirectly^7–13^ via paracrine signaling, to regulate glucagon secretion. At a more fundamental level, it is debated whether rising glucose levels stimulate^14^, suppress^15,16^, or do not affect^17^ glucagon secretion from α-cells. A recent study employing α-cell-specific GCaMP-expressing mouse models demonstrated heterogeneous patterns of glucose-stimulated cytosolic Ca^2+^ responses and glucagon secretion across α-cell subpopulations^18^. This aligns with our previous findings^5^, as well as those of others^19^, demonstrating that pancreatic α-cells exhibit both functional^5,19^ and genetic heterogeneity^19,20^. However, the mechanisms and physiological relevance of this heterogeneity remain unclear, particularly in the context of disrupted intra-islet signaling.

α-, β-, and δ-cells are closely positioned within rodent and human islets^21^. This anatomical juxtaposition is believed to support local feedback loops that fine-tune islet hormone secretion and glucose regulation^22^. Indeed, the suppressive actions of β-, and δ-cells on α-cell activity are well established; β-cells secrete insulin, which suppresses α-cell activity via insulin receptor signaling^11,13,23^, and urocortin 3, which regulates δ-cell activity^24^, while δ-cells release somatostatin, which inhibits glucagon release by acting on somatostatin receptors^8,10,12,13,25^.

Given that a rise in intracellular Ca²⁺ is the primary trigger for hormone release in α-, β-, and δ- cells^5,26,27^, approaches that simultaneously monitoring dynamic Ca²⁺ oscillations within these cells offer a convenient proxy for assessing hormone secretion at the single-cell level, particularly when examining interactions between neighboring cells, such as β- and δ-cells, which are anatomically adjacent to α-cells and are likely to exert regulatory influences.

Men and women differ in their susceptibility to developing T1D^28^ and T2D^29^. While significant sex differences in glucose metabolism, glycemic control, and insulin sensitivity have prompted substantial research into sex-dependent β-cell function in diabetes^30,31^, data on potential heterogeneity of α-cell function from a sex-dimorphic perspective are unavailable. Given that sex-specific mechanisms in islet function may underlie differential susceptibility and metabolic outcomes in disease^32^, advancing our understanding of sexually-dimorphic α-cell functions holds considerable potential for developing more effective and potentially sex-specific strategies to correct hyperglucagonemia in patients with diabetes.

In the present study, we employed both sex of age-matched reporter mice expressing the genetically-coded Ca^2+^ sensor GCaMP6 selectively in α-cells^33^. High spatiotemporal resolution dual-Ca²⁺ indicator confocal imaging was performed in fresh pancreas slices, wherein a red-shifted Ca^2+^ indicator was further loaded. This approach enabled simultaneous monitoring of the Ca^2+^ oscillations in α-, β- and δ-cells. We aimed to investigate the α-cell heterogeneity through 1) characterizing, at the single-cell level, α-cell Ca^2+^ oscillation patterns in response to glucose; 2) assessing the Ca^2+^ oscillation patterns in α-cells in relation to the activities of neighboring β- and δ-cells. By selectively blocking insulin and somatostatin signaling pathways, we evaluated whether glucose regulates α-cell glucagon secretion via intrinsic glucose sensing mechanisms, paracrine interactions, or both; 3) lastly, addressing the contribution of sex dimorphism to α-cell glucose responsiveness with or without perturbed paracrine signaling.

## Material and methods

### Generation of GluCre:GCaMP6f^fl/fl^ (GluCre:GCaMP6f) mouse model

All animal procedures complied with the University of Montreal Animal Care and Use Committee (CIPA2022-10,040 CM21022GRs to GAR). Mice had free access to water, regular chow and were housed under a 12-hour light/dark cycle. GluCre:GCaMP6f mouse model (Figure 1A) was generated by crossing GluCre mice (obtained from Jackson Laboratory Strain #:030663) to mice that express GCaMP6f downstream of a LoxP-flanked STOP cassette (The Jackson Laboratory, stock no. 028865).

**Figure 1:**
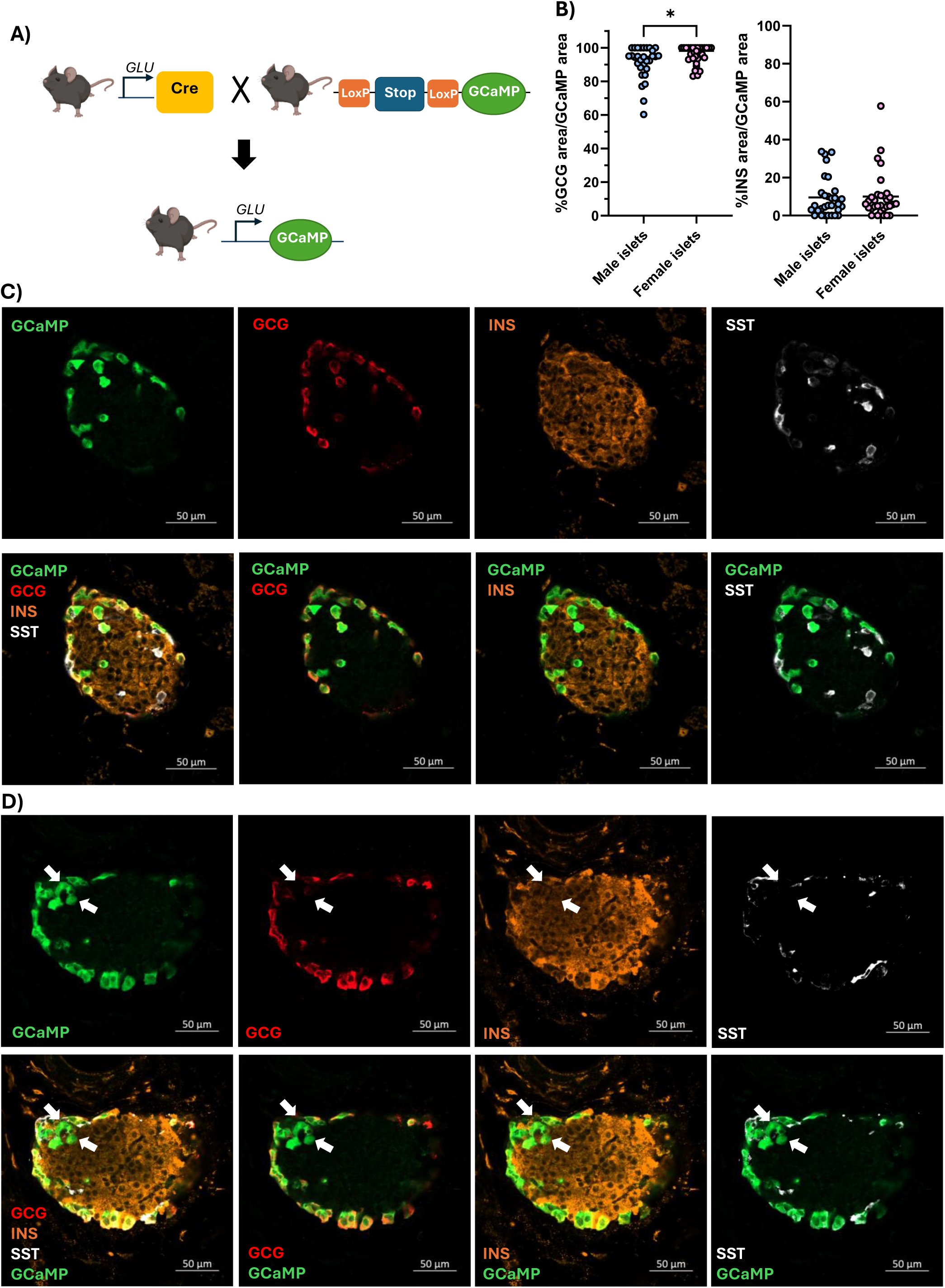
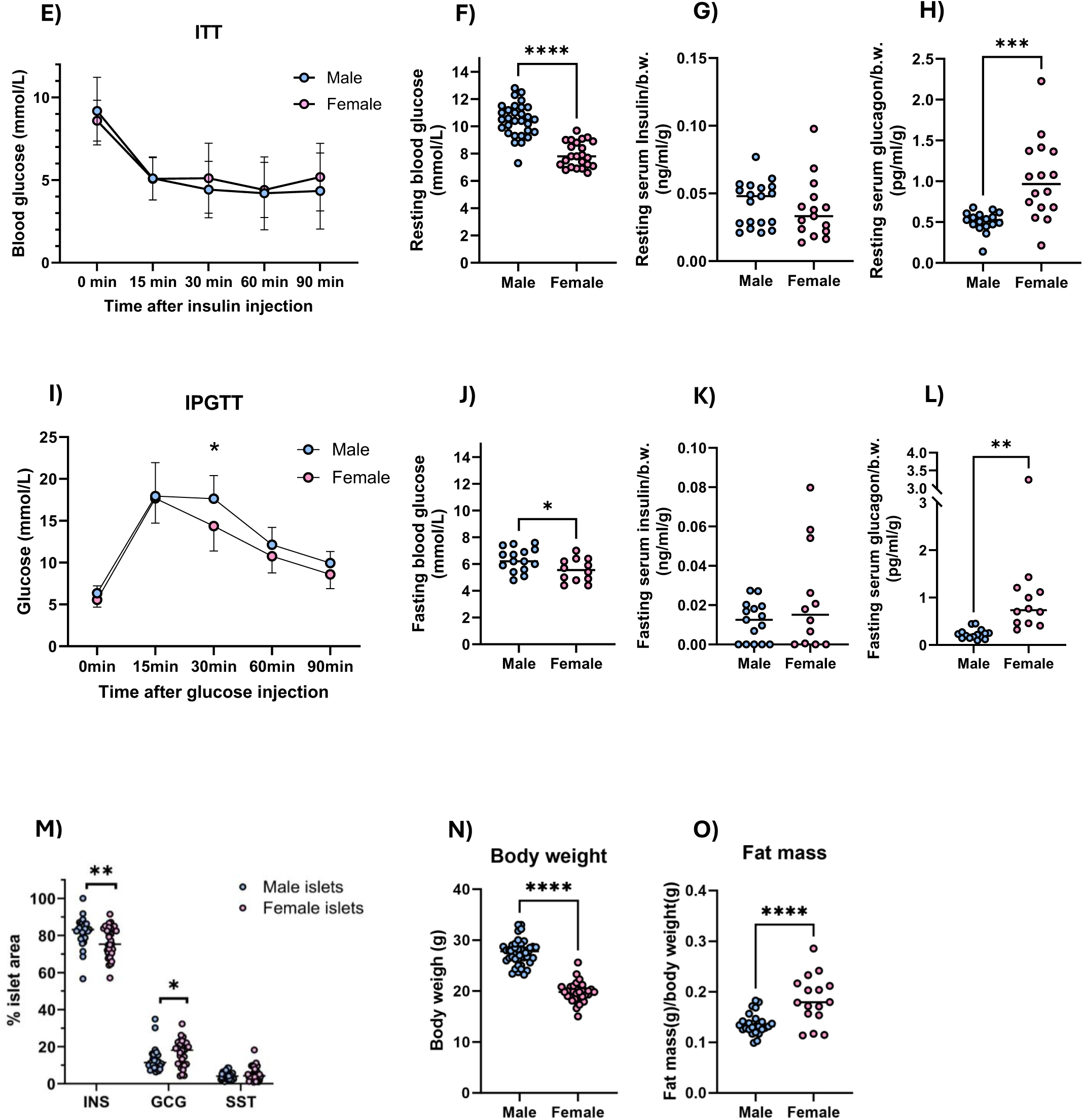
Characterization of GluCre:GCaMP6f^fl/fl^ mice. **(A)** Schematic diagram of GluCre:GCaMP6f^fl/fl^ mouse generation. Homozygous GluCre mice were crossed with homozygous GCaMP6f^fl/fl^ mice, in which GCaMP6f is expressed downstream of a LoxP-flanked STOP cassette. **(B)** Proportions of glucagon-positive (GCG, left) and insulin-positive (INS, right) areas within the total GCaMP-expressing area in individual islets from male (blue) and female (pink) mice. n = 30 islets from 6 males (76 ± 9.3 days), and 31 islets from 5 females (85.8 ± 7.8 days). *p < 0.05, unpaired two-tailed t-test. **(C-D)** Representative confocal images of male GluCre:GCaMP6f^fl/fl^ islets showing GCaMP expression (green) and immunostaining for glucagon (GCG, red), insulin (INS, orange), and somatostatin (SST, white). Arrows in **D)** indicate ectopic GCaMP expression in β-cells. **(E)** Insulin tolerance test (ITT). n = 15 males and 12 females (80.3 ± 12 and 82.8 ± 13.4 days, respectively). **(F)** Resting blood glucose levels. n = 30 males and 23 females (80.1 ± 1.2 and 85.8 ± 0.9 days). **(G-H)** Resting serum insulin **(G)** and glucagon **(H)** levels from 15 males and 12 females (79.3 ± 11.9 and 80.8 ± 13.4 days). ***p < 0.001, unpaired two-tailed t-test. **(I)** Intraperitoneal glucose tolerance test (IPGTT). n = 15 males and 12 females (84.3 ± 12 and 85.8 ± 13.4 days). *p < 0.05, two-way repeated measures ANOVA. **(J)** Fasting blood glucose levels from 15 males and 12 females (74.7 ± 8.2 and 82.7 ± 12.3 days). *p < 0.05, unpaired two-tailed t-test. **(K-L)** Fasting serum insulin (**I**) and glucagon (**J**) levels. Data represent 15 male and 12 female mice with mean age: 74.7 ± 8.2 and 82.7 ± 12.3 days, respectively. **p < 0.01, unpaired two- tailed t-test. **(M)** Quantification of INS-, GCG-, and SST-positive areas relative to total islet area. n = 30 islets (6 males, 76 ± 9.3 days) and 31 islets (5 females, 85.8 ± 7.8 days). *p < 0.05, **p < 0.01, Mann– Whitney test. **(N-O)** Phenotypic measurements including body weight (**N**) and fat mass (**O**) in 25 males and 16 females (86.2 ± 12.6 and 85.0 ± 13.8 days). ****p < 0.0001, unpaired two-tailed t-test. All animal age were provided as mean ± standard deviation.

### In vivo measurements

GluCre:GCaMP6f mice were either fasted for 16 hours (hrs) prior to fasting blood glucose measurement or assessed between 9:00-11:00 AM without fasting for resting blood glucose measurement. Blood glucose measurement, blood collection, intraperitoneal insulin (ITT) and glucose tolerance (IPGTT) tests were conducted as previously described^26,34^ or in Supplementary methods. Serum insulin and glucagon levels were measured using ELISA Kits (ALPCO Diagnostics, New Hampshire, USA; Mercodia, Uppsala, Sweden) per manufacturer’s instructions.

### Pancreas slice preparation, Ca^2+^ indicator loading, and immunoassay

GluCre:GCaMP6f mouse pancreas slices were prepared following protocols described previously^35^ or in Supplementary Methods. Pancreatic slices, additionally loaded with Calbryte 590^AM^ (15µM; 2-hrs), were imaged on a confocal microscope (Zeiss LSM-900, “Airyscan”) equipped with an incubation system for temperature control (37°C), and a 20x/0.8 Plan achromat objective. Images were acquired at 6.25Hz in a single plane (256X256 pixels) without averaging, using the 488 and 561nm laser lines for GCaMP6f and Cabryte 590^AM^ excitation, respectively. During recording, pancreas slices were continuously perifused with HEPES-extracellular solution (ECS)^35^ at 1ml/min supplemented with incremental (1.8, 3.6, 5.4, 7.2, 9.0, 10.8mM) glucose concentration and pharmacological compounds as indicated. S961 (Novo Nordisk (Denmark) compound sharing program), insulin (Novolin®ge Toronto, Novo Nordisk, ON, Canada), somatostatin-14 (Sigma-Aldrich, ON, Canada), MK4256 (MedChemExpress LLC, NJ, USA) and CYN154806 (Tocris Bioscience) were freshly prepared before each experiment. Movies were subjected to semi-automatized analytical pipeline that were developed previously^35,36^. After imaging, pancreas slices were immunoassayed following protocols^35^ described in Supplementary Methods.

### Classification of α-cells based on Ca^2+^ activity patterns

α-cells were categorized semi-automatically based on their Ca^2+^ activity patterns during glucose concentration ramps. We applied a custom Python script that segmented the recording into 6 time-intervals matching the six glucose concentrations on the ramp and quantifies Ca^2+^ events per segment per α-cell ROIs. For each cell, we calculated the spikes (events) per interval, total event count, and the relative distribution across intervals. Cells were categorized as: **“inactive”**, if all six time intervals exhibited fewer than five Ca^2+^ events; **“constantly active”**, if events were distributed relatively uniformly across all intervals, with a variation ≤15% between segments; **“low-glucose sensitive”**, if the combined activity during the first three intervals (corresponding to the low-glucose phase) exceeded that of the last three; and **“high-glucose sensitive”**, if the opposite pattern was observed, with greater activity in the final three intervals (high-glucose phase).

### Spatial and functional correlation analysis of α- and β-cells

Ca^2+^ oscillation traces from α- and β-cell ROIs within each islet were extracted and concatenated for analysis. Pairwise Pearson correlation coefficients were computed (detailed in Supplementary methods) between all α- and β-cell traces to quantify their functional relationships. For spatial analysis, the x-y coordinates of each ROI were used to calculate the distance between cell pairs. To investigate local paracrine interactions, we considered α- and β-cell pairs that are within 25µm apart, given that β-cells average 10–13µm in diameter and mouse α-cells are smaller than β- cells^5^. Negative correlations (*r* < 0) between nearby α- and β-cell pairs were visualized by plotting cell locations as scatter color-coded coordinates with lines connecting the negatively correlated pairs.

### Network analysis and cell active time

As described previously⁴³ and in Supplementary Methods.

### Statistics

Statistical analysis was performed (GraphPad Prism 10) with appropriate tests detailed in each figure legend; *p-value<0.05* was considered statistically significant.

### Data and code availability

Custom Python scripts developed to support the findings of this study are available on GitHub. For video processing: https://github.com/szarma/Physio_Ca/; for Ca^2+^ trace analysis: https://github.com/MSlakRupnik/Alpha_cell_analysis_for_publication. The raw data supporting the findings of this study are available from the corresponding authors upon reasonable request.

## Results

### *In vivo* characterization of the GluCre: GCaMP6f mouse

In this study, we aimed to uncover the dynamic behavior of α-cell Ca^2+^ activity across extensive cell populations through large-scale, high-resolution analysis. To this end, we generated GluCre:GCaMP6f mice which expressed GCaMP6, a Ca^2+^ sensor encoded downstream of the glucagon promoter, enabling GCaMP6 expression selectively in α-cells (Figure 1A).

Recombination in this model was highly efficient (Figure 1B), and quantitative immunocytochemical analysis revealed strong colocalization between GCaMP6 and glucagon-positive areas of the islet (Figure 1B-D); female GluCre:GCaMP6f mouse islets showed a greater degree of overlap between the glucagon positive areas and the GCaMP signal, compared to male islets (Figure 1B left and 1C; female vs. male: 90.4±2.24% vs. 87.4±2.58% of GCaMP areas are positive for glucagon). The remaining 10.1%±2.21% and 9.52%±1.93% of total GCaMP region in female and male islets, respectively, were positive for insulin, suggesting minor mis-expression of GCaMP6 in β-cells (Figure 1B right and 1D white arrows), as observed in a previous report^37^. For the present study, only cells that were positive for both GCaMP6 and glucagon were analysed as α-cells.

Insulin sensitivity and circulating fasting and resting insulin levels in GluCre:GCaMP6f mice were comparable between male and female mice (Figure 1E, 1G and 1K). However, the overall insulin-positive area within islets was smaller in female GluCre:GCaMP6f mice (Figure 1M), suggesting that β-cells in females exhibit greater secretory efficiency than those in males, thereby enabling comparable circulating insulin levels. Under an intraperitoneal glucose tolerance test (IPGTT), female mice exhibited better glucose clearance than males (Figure 1I). Female GluCre:GCaMP6f mice also showed lower fasting and resting glycemia (Figure 1F and 1J), lower body weight (Figure 1N), and higher body fat mass (Figure 1O), when compared to their age-matched male counterparts. Counterintuitively, female mice had higher circulating resting and fasting glucagon levels (Figure 1H and 1L), likely resulting from the larger glucagon-positive regions in female islets (Figure 1M).

### Glucose-stimulated Ca^2+^ dynamics in α-cells are heterogeneous

In order to explore intercellular interactions and Ca^2+^ dynamics in islets, we employed live pancreas tissue slice preparation and high spatiotemporal resolution confocal imaging^34,35^. A glucose concentration ramp incrementing from 1.8mM to 10.8mM over 60 minutes (Figure 2), mimicking changes in circulating glucose concentrations following a carbohydrate meal, was applied. In addition to recording the α-cells, which genetically expressing GCaMP6f, we further loaded a chemical Ca^2+^ indicator (Cabryte 590^AM^; emitting at 590nm; Figure 2, A-2) onto the slices to enable simultaneous tracking of Ca^2+^ activities in other cell types within the pancreas slices. Once recording was completed, the slices (140μm thick, ∼10-14 islet cell layers) were subjected to *post-hoc* immunoassay with anti-glucagon, -insulin and -somatostatin antibodies. For the scope of this study, we focused only on islet α- (GCaMP6- and glucagon-positive cells), β- and δ-cells (GCaMP6-negative, but Calbryte 590^AM^ -, insulin- or somatostatin-positive cells).

**Figure 2:**
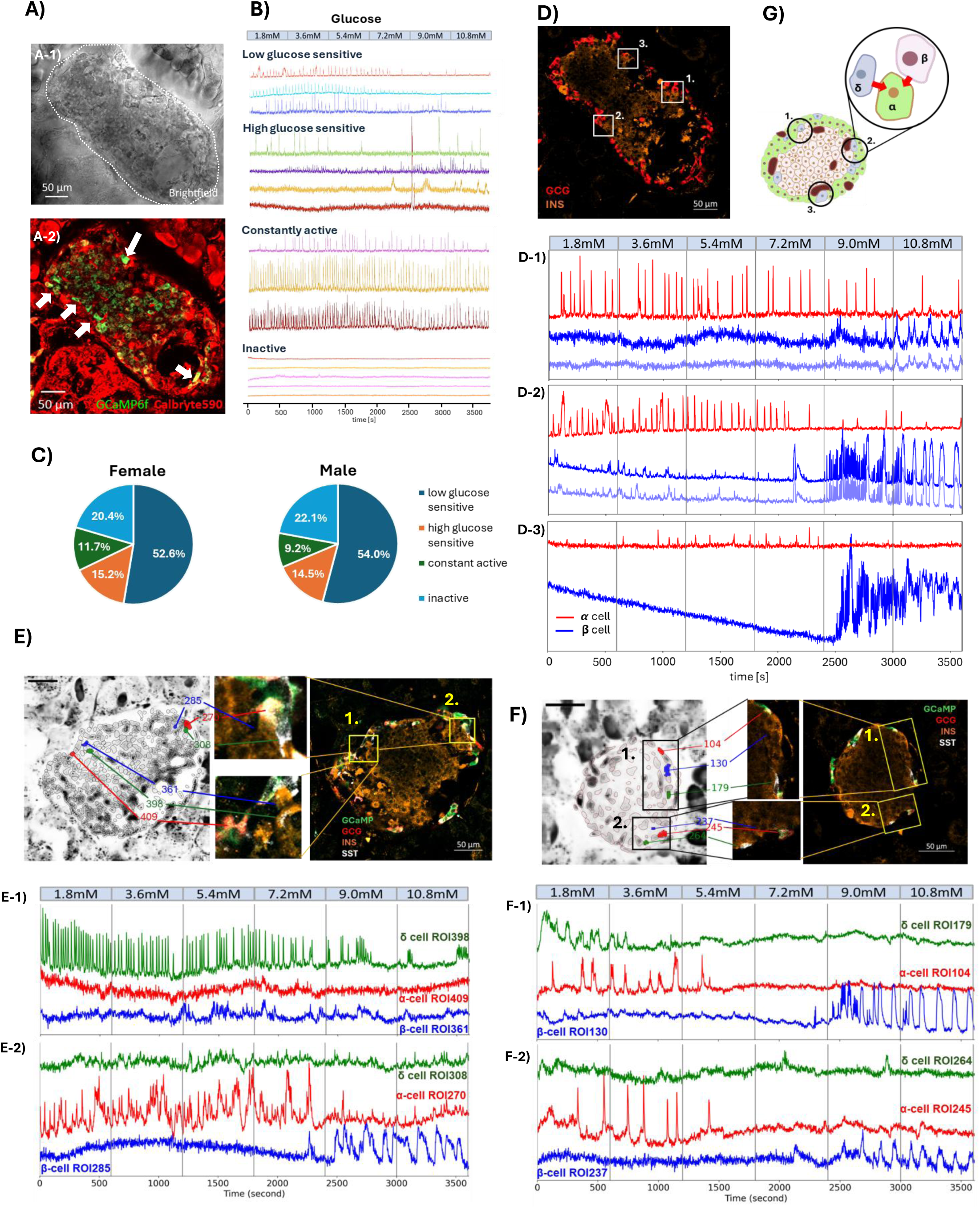
Ca^2+^ dynamics in α-, β- and δ-cells in GluCre:GCaMP6f^fl/fl^ islets in pancreas slice. **(A)** A representative islet within a pancreatic slice. **A-1)** Brightfield image of an islet (outlined by a white dotted line) in a pancreatic slice prepared from a G*luCre:GCaMP6f^fl/fl^* mouse. **A-2)** All pancreatic cells in the slice are loaded with Calbryte 590^AM^ (red), while α-cells express GCaMP6f (dense bright green; some examples are indicated by white arrows). Note the lighter green fluorescence in cells at the islet center represents inert signals, stemming from background fluorescence and no Ca^2+^ dynamics has emanated from these signals. **(B)** The pancreatic slice shown in (**A**) was perifused with an incremental glucose concentration ramp (top). Representative Ca²⁺ activity traces from α-cells in response to the glucose ramp are shown. These responses can be categorized into four groups: low glucose-sensitive, high glucose-sensitive, constantly active, and inactive α-cells. **(C)** Pie chart representations showing the proportions of types of α-cell response in islets in female (left) and male (right) mice. Data were collected from pancreas slices of 9 male and 5 female mice, containing 47 male and 29 female islets, respectively. A total of 2030 ROIs from male islets and 1289 ROIs from female islets were analyzed. Each ROI co-expresses GCaMP and glucagon. Every ROI is categorized according to their response to the incrementing glucose concentration ramp. The total number of ROIs in each category was then normalized to the total number of ROIs to derive the percentage proportion of each categorization. **(D)** *Post-hoc* immunostaining of the pancreatic slice shown in **(A)**, stained for glucagon (GCG, red) and insulin (INS, orange). **D-1)** Ca^2+^ activity profiles of α-cells (red) and β-cells (blue) corresponding to the inset white square in **(D)** labeled “**1.**”. The same applies to **D-2** and **D-3**, which correspond to inset squares labeled “**2.**” and “**3**.”, respectively. Note the heterogeneous Ca²⁺ responses of α-cells across different regions of the islet, which occur in anti-phase to the Ca²⁺ activity of neighboring β-cells within that particular region. **(E)** The top left image shows topologically defined regions of interest (ROIs) derived from a Ca²⁺ recording. The top right image shows the corresponding *post-hoc* immunostaining of the recorded islet, with GCaMP-expressing cells (green) and staining for glucagon (GCG, red), insulin (INS, orange), and somatostatin (SST, white). Inset squares **1.** and **2.** are enlarged in the top-middle panel, highlighting islet regions where α-, β-, and δ-cell ROIs are identified (left image) and confirmed by immunostaining (right image). The ROIs are numbered, and their corresponding Ca²⁺ activity profiles are shown in **E-1** and **E-2**. Note in **E-1:** an inactive α-cells is neighbored by an active δ-cells, whereas in E-2: an active α-cells is neighbored by a less active δ-cells. **(F)** This panel follows the same organizational layout as panel **(E)**, with regions of the islet (Insets 1 and 2) where α-, β-, and δ-cell ROIs are identified (left) according to the corresponding immunostaining (right), along with their respective Ca^2+^ activity profiles shown in **F-1** and **F-2**, respectively.

We observed that during an ascending glucose concentration ramp, Ca^2+^ dynamics in α-cells were markedly heterogeneous (Figure 2B). Whilst a majority of α-cells (52.6±4.2% and 54.0±3.1% in female and male islets, respectively; Figure 2B-Low glucose sensitive and Figure 2C) displayed robust Ca^2+^ firing activities at the low glucose range between 1.8mM to 5.4mM, a minority of α- cells responded to higher glucose concentrations (15.2±2.3% and 14.5±2.6% in female and male, respectively; Figure 2B-High glucose sensitive and Figure 2C)), or remained constantly active throughout the glucose ramp (11.7±1.9% and 9.2±1.5% of all α-cells in female and male, respectively; Figure 2B-Constantly active and Figure 2C). Approximately a quarter of α-cells were relatively quiescent throughout the entire glucose concentration ramp (20.4±2.6% and 22.1±2.3% in female and male islets, respectively; Figure 2B-inactive and Figure 2C)). We note that these were live α-cells, as evidenced by occasional spontaneous Ca^2+^ oscillations and their intrinsic GCaMP6 fluorescence, which indicated their viability despite being otherwise functionally silent. We defined cells exhibiting less than five spikes throughout the course of glucose ramp as inactive cells.

### Glucose-regulated Ca^2+^ dynamics in α-cells are associated with Ca^2+^ dynamics in neighboring β- and δ-cells

To explore a possible role for neighboring cells in controlling α-cell dynamics, we investigated α- cells, along with adjacent β- and δ-cells, across different topological locations within individual islets. We found that the Ca^2+^ activities in α-cells at different islet regions (Figure 2D) exhibited asynchronous patterns of Ca^2+^ activities between one another. Unlike β-cells^38^, α-cells exhibited weak connectivity in islets from both sexes (mean degree <1, clustering coefficient <0.15; Supplementary Figure 1). Remarkably, Ca^2+^ dynamics in α-cells were always in antiphase with the nearby insulin-positive β-cells (Figure 2D-1 to Figure 2D-3), aligned with earlier findings^39^. Pearson correlation analysis performed on juxtaposed α- and β-cells also revealed a negative correlation in their Ca^2+^ traces, confirming an antiphase relationship between the two cell types (Supplementary Figure. 2A, B). This pattern was further supported by Pairwise Pearson Correlation analysis of concatenated fluorescence traces from all α-cell and β-cell ROIs within individual islets (Supplementary Figure 2C).

**Figure 3:**
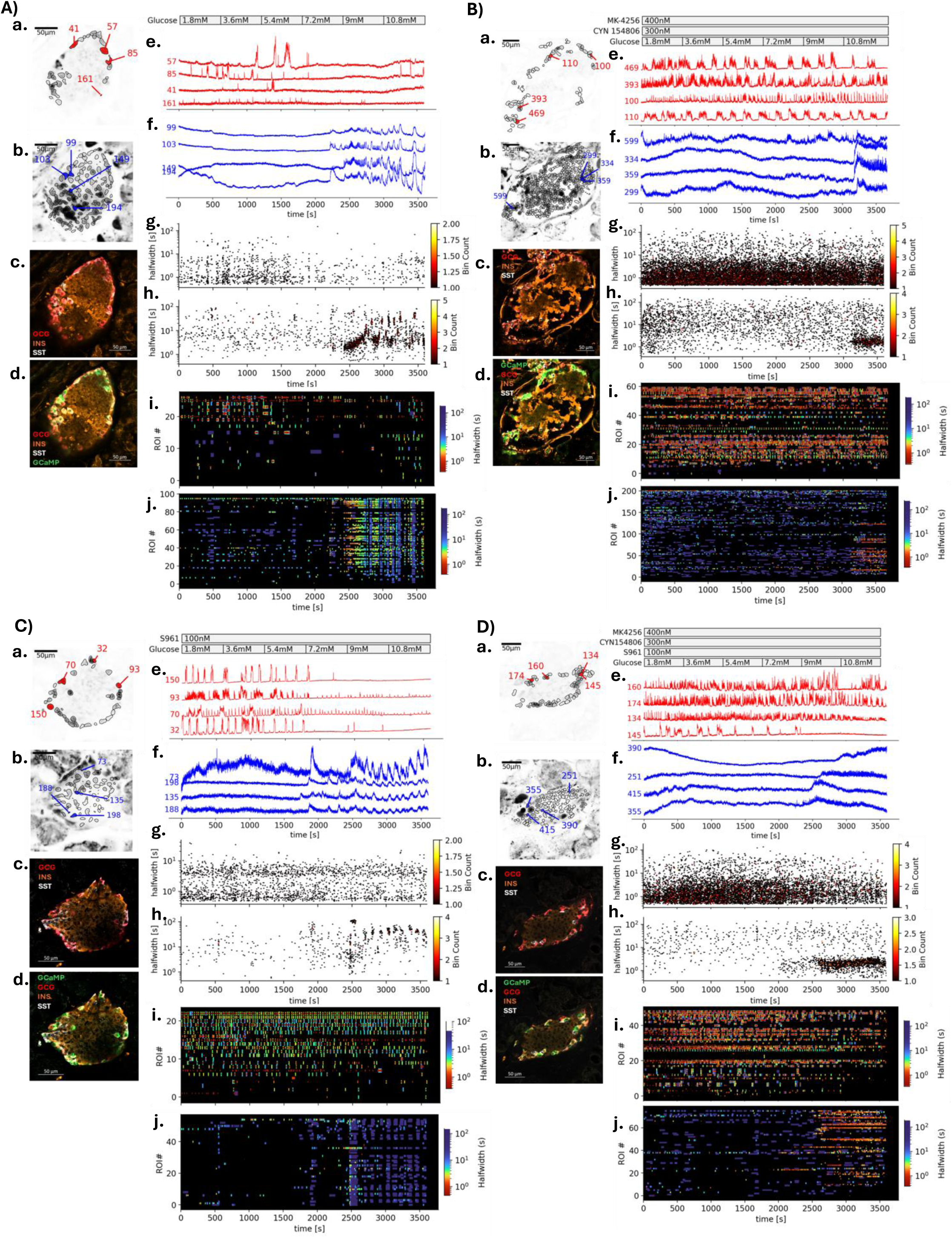
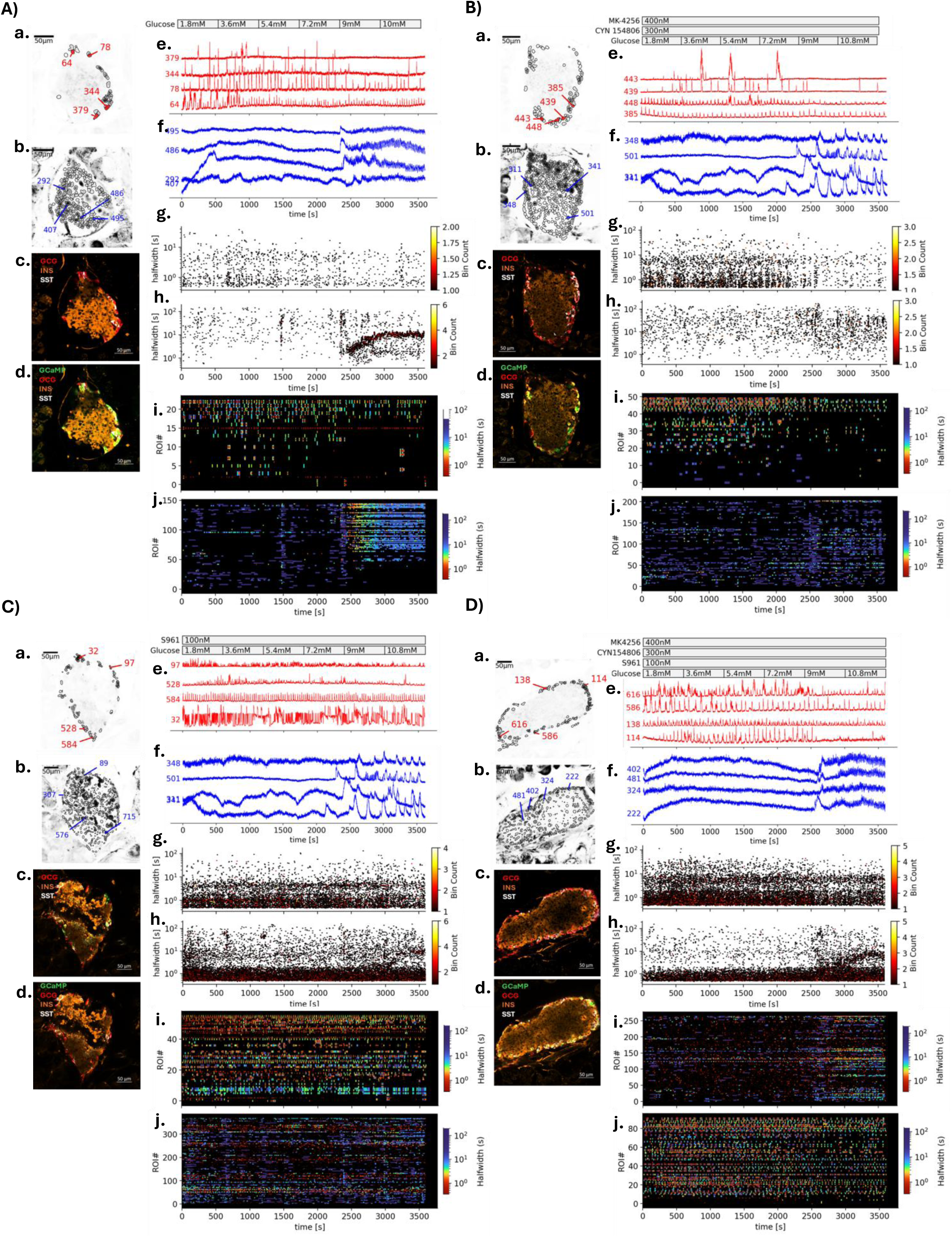
Modulation of α- and β-cell Ca^2+^ dynamics by insulin and somatostatin receptor antagonists in female and male mouse islets. **3-1 : (A)** Control condition. A representative female mouse islet exposed to an incremental glucose concentration ramp (top right). Regions of interest (ROIs) corresponding to α- and β- cells (**a.** and **b.**, respectively) are shown in the left panel. Immunostaining for glucagon (GCG), insulin (INS), and somatostatin (SST) **(c.)**, as well as GCaMP expression (**d.**), is displayed. Panels **e.** and **f.** show representative Ca^2+^ dynamics of the α- and β-cells labeled in **a.** and **b.** Panels **g.** and **h.** show hexbin plots representing Ca^2+^ transient distributions in all α- (**g.**) and β- cell (**h.**) ROIs from this islet. Events are plotted as a function of peak time (x-axis; time [s]) and duration (y-axis; halfwidth [s]). Color intensity indicates event density according to the color bar on the right. Panels **i.** and **j.** are raster plots of Ca^2+^ transients in α- (**i.**) and β-cell ROIs (**j.**), respectively, with each dot representing an event and color indicating event duration. An adjacent color bar (right) shows the mapping of halfwidth values on a logarithmic scale. **(B–D)** As in **(A)**, panels (**a. to j.**) show islet morphology, calcium dynamics, and event distributions under pharmacological modulation during the glucose ramp. **(B)** Insulin receptor antagonist (S961), **(C)** somatostatin receptor antagonists (MK-4256 and CYN154806), and **(D)** combined treatment (S961, MK-4256 and CYN154806) were applied throughout the ramp to assess their effect on Ca^2+^ activity in α-and β-cells. **3-2** replicates layout of Figure 3**-1** **(A–D, with subpanels a–j)** for male mouse islets, allowing direct comparison of treatment effects between sexes.

We repeatedly observed antiphase activity between juxtaposed α- and β-cells in all islets examined. This suggested that either direct interactions exist between α- and β-cells, or that a third cell type (i.e., δ-cells) drives the “on-and-off” activity between α- and β-cells. Hence, we examined the Ca²⁺ dynamics in δ-cells that were neighboring α-/β-cell pairs (Figure 2, E-F). An inverse relationship in Ca²⁺ activity was observed between δ-cells and adjacent α-cells during ramp stimulation, particularly within the glucose range between 1.8–7.2mM when β-cells were largely inactive. We observed that active δ-cells were frequently adjacent to less active α-cells (Figure 2E-1), and conversely, active α-cells were often juxtaposed to less active δ-cells (Figure 2E-2). This spatial relationship was also evident in temporal activity patterns: as δ-cell Ca²⁺ activity declined (1.8–5.4mM glucose; Figure 2F-1), the nearby α-cell activity often increased. Conversely, periods of low δ-cell activity frequently coincided with heightened α-cell activity (1.8–5.4mM glucose; Figure 2F-2), with δ-cell activity resuming (7.2–9.0mM glucose; Figure 2F-2) as α-cell activity declined. Quantification of active time in these cell pairs supported an inverse relationship in the robustness of Ca^2+^ oscillations between juxtaposed α- and δ-cells (Supplementary Figure 3A). Of note, no obvious association was observed between δ- and the neighboring β-cell Ca²⁺ dynamics (Figure 2E-1, E-2 and Figure 2F-1, F-2) and the Ca^2+^activity in δ-cells was notably heterogeneous (Supplementary Figure 3B). It was therefore difficult to define a general pattern of δ-cell Ca^2+^ activities in response to the incremental glucose ramp, as responses varied, showing increases, decreases, or no change over the courses of ramp stimulation. Nevertheless, our observations indicated the possibility that α-cells are under the control of both β-cell (anti-phase Ca^2+^ activity relationship between α- and β-cells) and δ-cells (inverse relationship in the strength of Ca²⁺ activity between α- and δ-cells).

### Somatostatin, insulin, and glucose regulate α-cell Ca²⁺ activities

Within the islet microenvironment, α-cells in different regions are surrounded by differing numbers of β- and δ-cells (Figure 2G, islet regions 1-3). Depending on the precise cell type composition, each of these different combinations could theoretically provide differential inputs into the nearby α-cells (Figure 2G, inset circle red arrows), given that insulin and somatostatin, secreted by β- and δ-cells respectively, are well-known inhibitors of α-cells^40^. We therefore hypothesized that the interaction of α-cells with different neighboring combinations of β- and δ- cells underlies, or may contribute to, α-cell functional heterogeneity. Accordingly, administering either insulin receptor and/or somatostatin receptor antagonists during the glucose ramp on pancreas slices should shift most of α-cell Ca²⁺ activity toward hyperactivity, thereby altering heterogeneity.

### δ-cell control of α-cell function

We first tested paracrine regulation of α-cell Ca²⁺ activity by δ-cells by using either somatostatin or <u>s</u>omato<u>s</u>tatin receptor <u>a</u>ntagonists (SSTRA: 400nM MK4256 and 300nM CYN154806, which antagonize somatostatin receptors type 2 and 3 that are expressed mainly in mouse islet α- cells^41,42^). At a steady glucose level of 3.6 mM, where α-cell Ca²⁺ activity was high, SST inhibited α-cell Ca^2+^ activity in a dose-dependent manner, with stronger inhibition observed in α- cells from female islets (Supplementary Figure 4 A-B inset). Conversely, SSTRAs potentiated α- cell Ca^2+^ spike frequency in female islets (Supplementary Figure 4D) but did so only in few male islets (Supplementary Figure 4D). These results indicate that female α-cells are more sensitive to δ-cell–mediated somatostatin inhibition.

During glucose ramp stimulation, SSTRA triggered significant increases in α-cell Ca^2+^ activity in both female and male islets (Figure 3-1B; 3-2B; Figure 4 top panel: SSTRA treatment).

**Figure 4:**
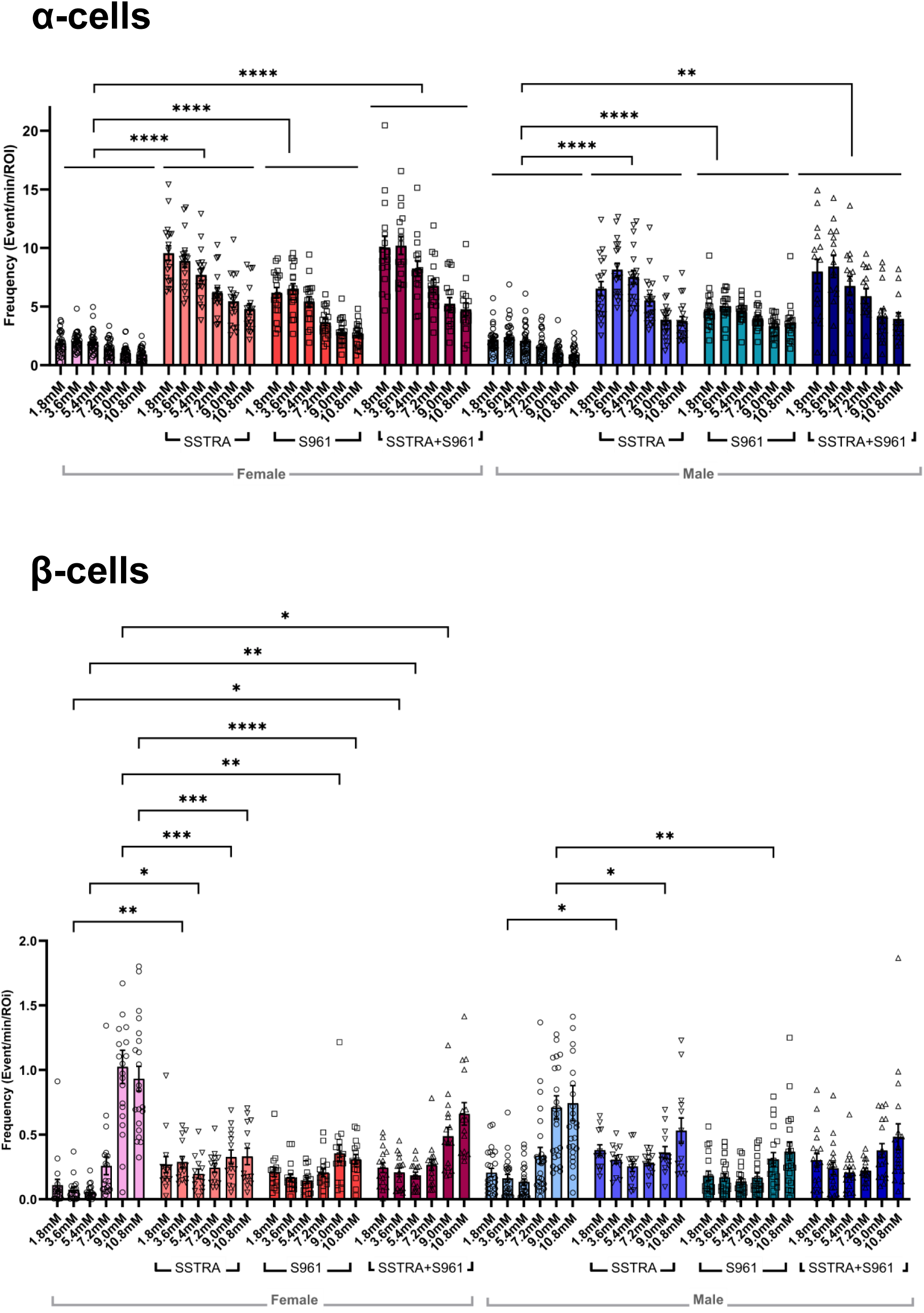
Summary of α- and β-Cell Ca^2+^ activity response across different glucose concentrations and receptor antagonists. Ca²⁺ oscillation frequencies in α-cells (top panel) and β-cells (bottom panel) were measured in response to an increasing glucose ramp alone or in combination with receptor antagonists. Bars represent mean ± SEM Ca²⁺ oscillation frequencies in female (pink, orange, red, dark red) and male (light blue, purple, green, navy) islets under the following conditions: glucose ramp alone; glucose ramp + SSTR antagonists (SSTRA: 400 nM MK4256 + 300 nM CYN154806); glucose ramp + insulin receptor antagonist (S961: 100 nM); or glucose ramp + SSTRA + S961. Each data point, represented by empty circles, downward triangles, squares, or upward triangles, indicates the mean Ca^2+^ activity frequency of α-cells (top) or β-cells (bottom) of an islet. **Top panel α-cells:** Glucose ramp only: 30 female and 31 male islets from 5 mice per sex (females, 85 ± 7.8 days; males, 76 ± 9.3 days); SSTRA: 18 female and 19 male islets from 3 mice per sex (females, 65.3 ± 5.5 days; males, 77.3 ± 3.8 days); S961: 19 female and 19 male islets from 3 mice per sex (females, 84 ± 2.0 days; males, 86.5 ± 10.1 days); SSTRA + S961: 16 female and 15 male islets from 3 mice per sex (females, 88 ± 7.0 days; males, 94.3 ± 6.4 days). **Bottom panel β-cells:** Glucose ramp only: 21 female and 25 male islets from 5 female and 7 male mice (females, 85 ± 7.8 days; males, 78.7 ± 11.2 days); SSTRA: 14 female and 13 male islets from 3 mice per sex (females, 65.3 ± 5.5 days; males, 77.3 ± 3.8 days); S961: 16 female and 19 male islets from 3 mice per sex (females, 84 ± 2.0 days; males, 86.5 ± 10.1 days); SSTRA + S961: 16 female and 18 male islets from 3 mice per sex (females, 88 ± 7.0 days; males, 94.3 ± 6.4 days). Mix-effects model analysis was performed; Significance levels are indicated as: *p* < 0.05 (\**), p < 0.01 (**), p < 0.001 (\*\*\**), and *p* < 0.0001 (****).

Quantitative analysis of α-cell Ca²⁺ activity in individual islets showed that SSTRA treatment increased the proportion of constantly active α-cells (Figure 5A-B: 37.4±5.5% in females, 21.9±3.3% in males, *p*<0.0001 in females), more potently in the female islets (*p*<0.05; Figure 5D). The proportion of inactive (8.3±1.6% in females, 17.2±3.4% in males; Figure 5A and B) and high glucose-sensitive α-cells under SSTRA treatment remained otherwise comparable to those observed with glucose ramp alone. (13.5±2.3% in females, 9.6±2.7% in males; Figure 5A-B).

**Figure 5:**
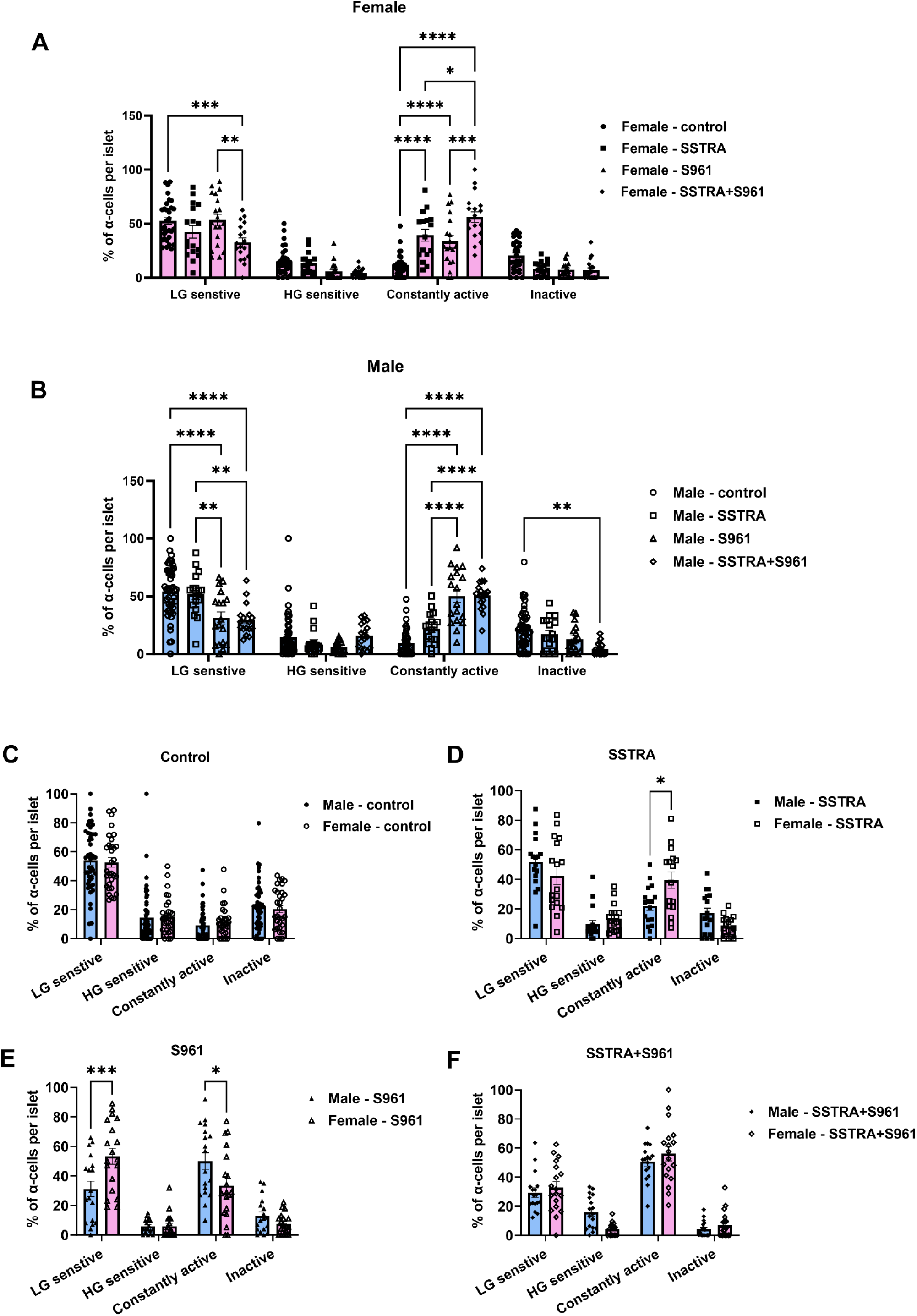
Classification of α-cell Ca²⁺ activity profiles in female (A) and male (B) islets across treatments. Islets from Female (**A**) and male (**B**) pancreas slices treated with: glucose ramp alone (control), or in combination with SSTRA, S961, or both. α-cells in each islets were categorized based on their Ca²⁺ oscillation profiles into four groups as defined in Figure 2C: low glucose (LG) sensitive, high glucose (HG) sensitive, constantly active, or inactive. **C-F:** Sex-based comparison of α-cell proportions in each oscillation category: glucose ramp alone (control; **C**), +SSTRA (**D**), +S961 (**E**), or SSTRA+S961 (**F**). Each symbol represents the percentage of α-cells relative to the total α-cell population per islet; each bars indicates mean ± SEM. Two-way ANOVA with Tukey’s multiple comparisons test was performed; Significance levels are indicated as: *p* < 0.05 (\**), p < 0.01 (**), p < 0.001 (\*\*\**), and *p* < 0.0001 (****). Female islets, data were collected from 29 islets (5 mice, 85 ± 7.8 days old) under glucose ramp alone; 17 islets (3 mice, 77.3 ± 3.8 days) with SSTRA; 19 islets (3 mice, 86.5 ± 10.1 days) with S961; and 18 islets (3 mice, 94.3 ± 6.4 days) with SSTRA + S961. Male islets: glucose ramp alone included 47 islets (9 mice, 78.7 ± 11.2 days); SSTRA included 17 islets (3 mice, 65.3 ± 5.5 days); S961 included 19 islets (3 mice, 84 ± 2 days); and SSTRA + S961 included 6 islets (3 mice, 88 ± 7 days).

These findings suggest that previously inactive α-cells may have transitioned into a constantly active state as a result of SST receptor antagonism. The notion that inhibition by somatostatin, presumably mediated by δ-cells under physiological conditions, contributes, at least to some extent, to the apparent heterogeneity of α-cell Ca²⁺ activity is supported.

Of note, during the SSTRA-supplemented glucose ramp, β-cell Ca²⁺ activity frequency was increased at basal glucose concentrations (1.8-5.4mM glucose in both sex; Figure 4 β-cells: SSTRA treatment). However, the decline in α-cell activities and the onset of synchronized β-cell Ca²⁺ activities, particularly in female islets (Figure 3-1B: g-j; raster plots), were both right-shifted towards higher glucose concentrations. That is, in the presence of SSTRA, α-cells remain activated above 5.4mM glucose, where Ca²⁺ oscillations in most of α-cell typically declines, whereas β-cells showed a significantly blunted Ca²⁺ oscillation frequency at ≥ 9mM glucose in islets of both sex (Figure 4 β-cells: SSTRA treatment), as a result of the delayed onset of their synchronized Ca^2+^ activation during the glucose ramp (Figure 3-1B; Figure 6A). This is also evident in the Ca²⁺ activity density plot (Supplementary Figure 6C), where α-cell Ca²⁺ activity from all islets under SSTRA treatment is compiled.

**Figure 6:**
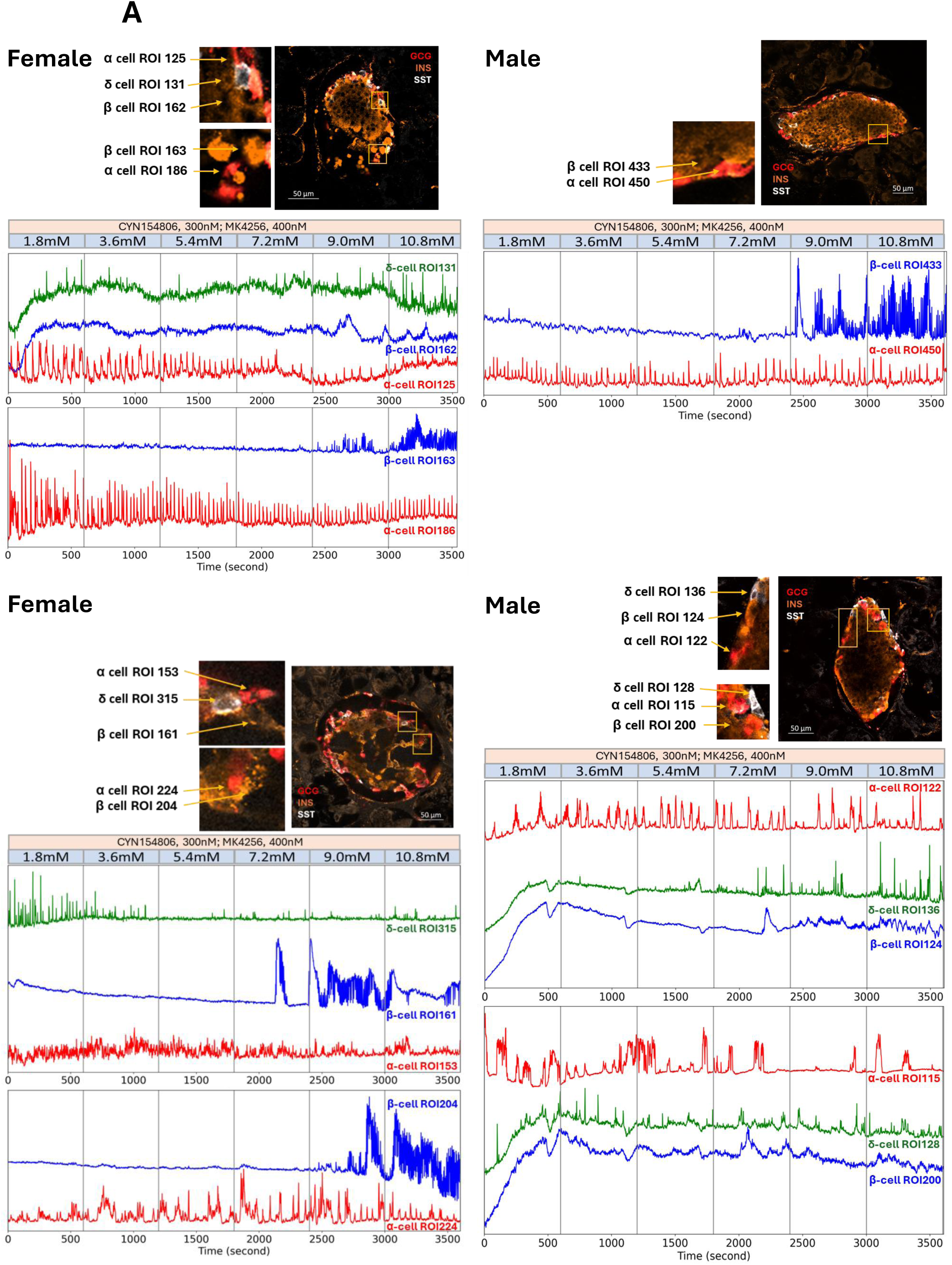
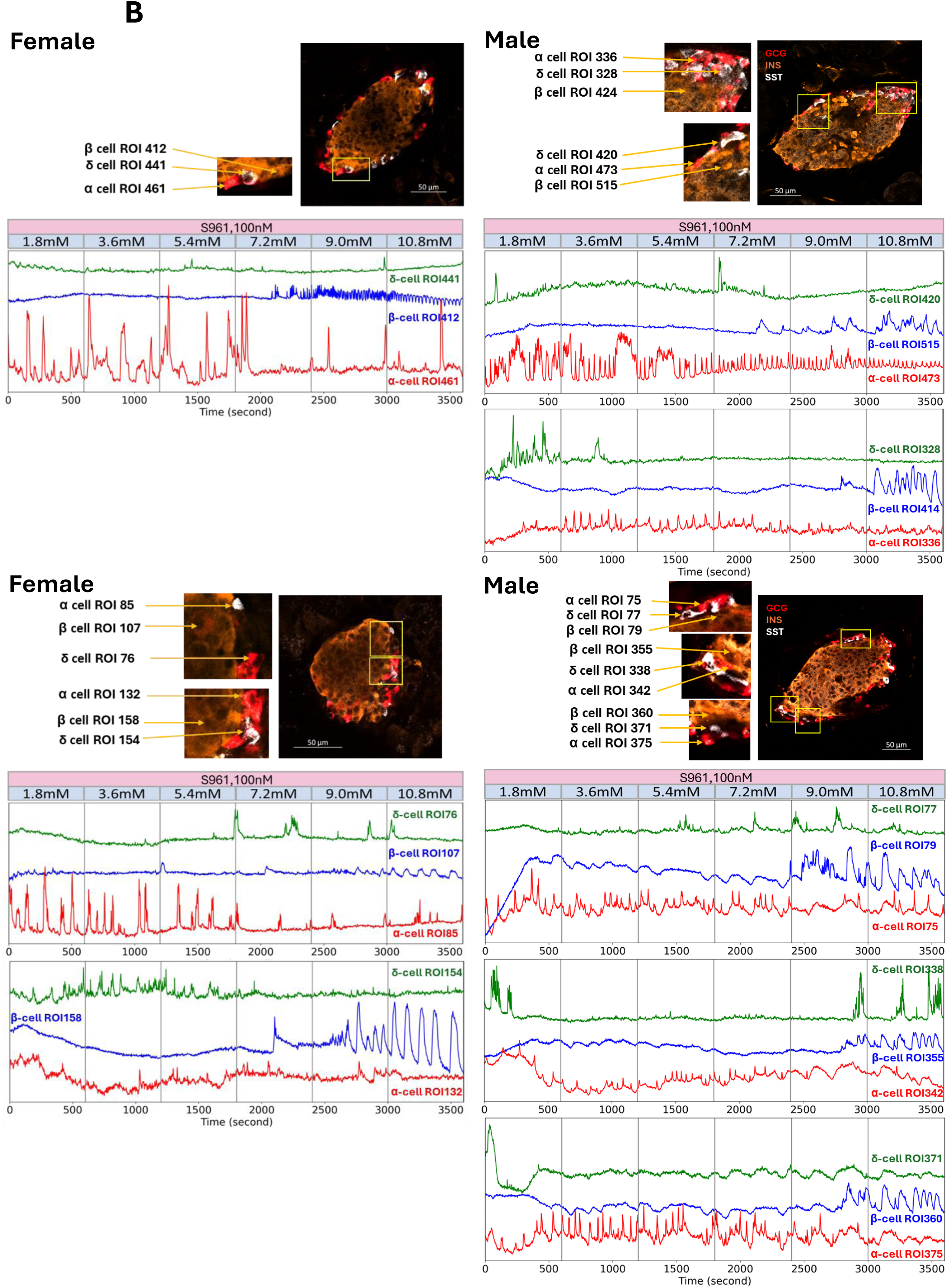
δ cell Ca²⁺ oscillation profiles in relation to neighboring β and α cells under SSTRA (6A) and S961 (6B) treatments. **6A** and **6B** shared the same organizational layout. The left side of the figure displays two representtive female islets, while the right side shows two male islets, each stained for insulin (INS, orange), glucagon (GCG, red), and somatostatin (SST, white). Yellow squares within the islet images highlight regions containing identifiable α-, β-, and δ- cells, which are enlarged to the left of the islet images. Below these images, Ca^2+^ oscillation profiles of δ- (green), β (blue), and α-cells (red) within the yellow squares are displayed and labeled with their respective regions of interest (ROI). Note in **6A,** the Ca^2+^ oscillations between δ-cells (green traces) and α-cells (red traces) no longer display an inverse relationship. In contrast, in **6B**, this inverse relationship between δ-cell and α-cell oscillations is still present.

Given that a chemical Ca²⁺ indicator was loaded into each slice, we explored whether δ-cell activity could be evaluated during SSTR agonist treatment. We found that δ-cell Ca²⁺ activity intensity relative to α-cell Ca²⁺ activity no longer exhibited the reciprocal pattern observed in the absence of the inhibitor (Figure 6A).

### β-cell control of α-cell function

We next tested the paracrine effect of β- on α-cells using insulin or the insulin receptor antagonist, S961^43^. At 3.6mM glucose, despite the addition of insulin, α-cell Ca^2+^ activity was only modestly reduced (Supplementary Figure 5A-B). Consistently, the insulin receptor antagonist S961 did not significantly potentiate Ca^2+^ activity in α-cells in both female and male pancreas slices (Supplementary Figure 5C-D). We therefore preincubated pancreas slices with S961 for 1hr before subjecting them to a S961-supplemented glucose ramp. During the glucose ramps, S961 significantly enhanced α-cell Ca²⁺ activity over low glucose ranges (Figure 3-1C, 3- 2C, and Figure 4 top: S961 treatment). Particularly in male islets, S961 treatment notably altered α-cell glucose sensitivity (Figure 3-2C; Supplementary Figure 6D), shifting the suppression of α-cell Ca²⁺ oscillations and the activation of synchronized β-cell Ca²⁺ oscillations to occur at higher glucose concentrations (>7.2mM). Consistent with this sex-dependent sensitivity of S961, quantitative analysis of α-cell Ca²⁺ activity revealed that S961 treatment increased the proportion of constantly active α-cells to 33.6±5.4% in females vs. 48.1±5.6% in males (*p*<0.0001 comparing to control; Figure 5A and 5B), with higher potency in male islets (*p<*0.05; Figure 5E), whereas the proportion of inactive (7.5±1.6% in females, 13.2±2.6% in males; Figure 5A and 5B) and high glucose-sensitive α-cells (5.7±1.9% in females, 5.5±1.1% in males; Figure 5A and 5B) were modestly reduced. These results reinforced the hypothesis that insulin inhibition, presumably mediated by β-cells in physiological contexts, also contributes, at least in part, to the heterogeneity of α-cell Ca²⁺ activity.

Similar to SSTRA treatment, β-cell Ca^2+^ oscillation frequencies under S961 treatment were blunted at ≥9mM glucose (Figure 4 bottom panel, S961 treatment). Moreover, β-cell activity under S961 treatment was no longer strictly in antiphase with that of α-cells when both cell types were active (Figure 6B). Assessment of δ-cell Ca²⁺ activity under S961 treatment revealed that the intensities of α- and δ-cell Ca²⁺ activities continued to exhibit an inverse pattern (Figure 6B).

### Effects of glucose on α-cell Ca^2+^ activity

Lastly, we applied SSTR antagonists and S961 together (Figure 3-1D and 3-2D). The combined antagonism of somatostatin and insulin receptors resulted in about half of α-cells shifting to a constantly active state (56.1±4.9% in females, 50.6±3.1% in males; *p*<0.0001 for both sexes; Figure 5A-B: SSTRA+S961), reducing the proportion of inactive α-cells (6.9±2.2% in females, 4.2±1.3% in males; *p*<0.01 in males; Figure 5A-B). Meanwhile, only approximately 30% of α- cells remained sensitive to low glucose (32.9±4.0% in females, 29.3±3.3% in males; both *p*<0.0001 vs. control; Figure 5A-B SSTRA+S961). Consequently, the overall α-cell Ca²⁺ activities, although significantly increased (Figure 4 top panel-SSTRA+S961), were still glucose concentration-dependent (Figure 4, top panel-SSTRA+S961; supplementary Figure 6B).

## Discussion

In this study, we demonstrated that during incremental glucose ramps, α-cell Ca²⁺ oscillations exhibited marked heterogeneity. Importantly, α-cell Ca²⁺ oscillations occurred in anti-phase with neighboring β-cell Ca²⁺ oscillations and the intensity of α-cell Ca²⁺ oscillations were inversely associated with that of the neighboring δ-cell’s Ca²⁺ oscillations. We demonstrated that interfering with the paracrine communication, either between α- and β-cells and/or α- and δ-cells, by blocking either insulin receptor signaling alone, somatostatin receptor subtypes-2 and 3 alone, or both, led to large proportion of α-cells shifting toward hyperactivity. Interestingly, this occurred in a sex-dependent manner: female α-cells exhibited enhanced sensitivity to somatostatin-mediated signaling, whereas male α-cells were more responsive to insulin receptor signaling.

Collectively, we suggest that α-cell excitability is finely tuned by inhibitory signals arising from the close juxtaposition of insulin- and somatostatin-secreting β- and δ-cells, in addition to α-cells’ *intrinsic* glucose sensitivity, likely involving intracellular metabolism of glucose and regulation of ATP-sensitive K^+^, Na^+^ and Ca^2+^ channels^44^. We note that the latter mechanism presumably persists even when SSTR and InsR signalling are blocked (Figure 4), consistent with the activating effects of deleting the key metabolic glucose sensor, glucokinase, selectively in α- cells^45^. These findings highlight the critical role of intra-islet coordination between different cell types in maintaining α-cell responsiveness. They also suggest that sex differences in paracrine signaling may contribute to different glucagon secretion profiles in metabolic disease.

We demonstrated that suppression of SSTR2 and SSTR3 signaling leads to hyperactive α-cell Ca^2+^ oscillations, with sustained Ca^2+^ oscillation at high glucose concentrations (>5.4mM) in females. To date, there is no evidence for sex-dependent differences in SSTR2 and SSTR3 expression in the mouse α-cells. However, it has been suggested that the female sex hormone, estrogen, modulates SSTR expression in other tissues such as pituitary^46^, prostate^47^, and certain tumor cells^48^. It is therefore plausible that α-cells in female mice also express elevated levels of SSTR2 and SSTR3, owing to higher estrogen levels, leading to prolonged glucose-stimulated α- cells Ca^2+^ activity under SSTR2 and SSTRA3 antagonism.

We further revealed that insulin receptor antagonism by S961 leads to sustained α-cell Ca²⁺ activity in male mice under high glucose conditions, suggesting that male α-cells are more sensitive to insulin action. This heightened insulin sensitivity in male α-cells implies that, under conditions of insulin resistance in T2D or after β-cell loss in T1D, males may experience more pronounced hyperglucagonemia. This mechanism may help explain the higher prevalence of T1D or T2D in men^28,29^, as worsen hyperglucagonemia in male α-cell could drive excessive hepatic glucose production, contributing to both fasting and postprandial hyperglycemia.

We showed that, during an intraperitoneal glucose tolerance test (IPGTT), female GluCre:GCaMP6f mice exhibited better glucose clearance comparing to males despite having higher circulating glucagon levels. Females also had lower body weights compared to their male counterparts. These findings are consistent with previous reports demonstrating that chronic glucagon agonism^49^, or glucagon and GLP-1 co-agonism^50^, exerts antidiabetogenic effects in mice, including greater weight loss and improved glucose metabolism. Consistently, in a rat model, multiple low-dose glucagon injection into the mediobasal hypothalamus led to decreased hepatic glucose production and subsequent improved glucose tolerance^51^, mirroring our observations in IPGTT in female GluCre:GCaMP6f mice. These effects are likely because glucagon is capable of crossing the blood–brain barrier^52^ and bind to glucagon receptors in the hypothalamus and brainstem^53,54^, which are regions involved in regulating glucose metabolism, energy expenditure, and appetite. Glucagon also stimulates hepatic synthesis and release of fibroblast growth factor 21 (FGF21), a circulating peptide hormone that contributes to energy homeostasis via centrally mediated mechanisms^55^. Lastly, glucagon can elicit appetite-suppressing effect through the liver–vagus–hypothalamus axis^56^. Together, the central and peripheral actions of glucagon provide a mechanistic explanation for the improved glucose tolerance and reduced body weight observed in female GluCre:GCaMP6f mice in vivo.

We showed that islet insulin-positive area was smaller in female GluCre: GCaMP6f mouse islets (Figure 1M), yet insulin sensitivities and circulating insulin levels were comparable between female and male GluCre: GCaMP6f mice. This suggests that female β-cells are more efficient in producing insulin than their male counterparts. This observation aligns with previous research^30^, which demonstrated that wild-type female mouse islet β-cells exhibit greater insulin secretion efficiency, attributed to more depolarized glucose-induced membrane potential and increased firing frequency. Consistently, clinical studies found that women have higher insulin secretion for a given glucose load^57^. *Ex vivo* glucose-stimulated insulin secretion assays performed on cadaver donor human islets obtained from age-, BMI-, and HbA1-matched donors also showed greater insulin secretion in female islets compared with male islets, despite comparable β-cell content in islets between sex^58^.

We uncovered that δ-cells Ca^2+^ dynamics in response to an increasing glucose ramp are heterogeneous, with no obvious pattern, and are not strictly associated with the Ca^2+^ oscillation pattern of the neighboring β-cells. Conversely, the rigor of δ-cells Ca^2+^ oscillation is always in an inverse fashion to that of α-cells. These findings deviate from previous reports suggesting δ-cells Ca²⁺ spike increases with elevating glucose concentration^59^, as a result of direct electrical coupling and metabolic communication with between β- and δ-cells through gap junctions^8^. Since δ-cells have a neuron-like morphology characterized by a compact soma and dendrite-like extensions^59^, combined with their low abundance in islet and small cell size, we acknowledge that smaller sample size of δ-cells were successfully loaded, recorded, and confirmed as somatostatin-positive with *post-hoc* immunostaining in our study. Future studies employing mouse models such as SST:GCaMP with expanded δ-cell sample size will be instrumental in enabling more precise characterization of δ-cell Ca²⁺ dynamics across a range of glucose concentrations.

A recent study found that removing SST signaling in isolated mouse islets lowered the glucose threshold for β-cell Ca²⁺ oscillations^60^. In contrast, our work in fresh pancreas slices showed that SST signaling blockade prolongs α-cell hyperactivity and delays activation of synchronized β- cell Ca²⁺ oscillations, particularly in female islets, requiring higher glucose levels. We consistently observed this delay among β-cells when α-cell hyperactivity was induced, either by SST receptor antagonism in female islets or insulin receptor antagonism in male islets. These findings suggested that α- and β-cells coordinated closely to shape overall islet function, which is consistent with data presented herein and that of Ren et al^39^, showing antiphase (phase-locked) Ca²⁺ oscillations between α- and β-cells. It is therefore intuitive that prolonged α-cell activity delays the onset of synchronized β-cell oscillation during the incremental glucose ramp observed in this study. In the study by Huang et al^60^, only a subset of β-cells expressed GCaMP in the mouse model used, and α-cell Ca²⁺ dynamics were unfortunately not examined in parallel. These limitations hindered the mechanistic insight into why activation thresholds for synchronized β- cell activity are reset to lower glucose concentrations in response to SST signaling disruption, particularly given the tightly coordinated regulation between α- and β-cell function. Lastly, we note that differences in experimental models, i.e., acute pharmacological SST blockade in intact pancreas slices versus chronic SST depletion or genetic δ-cell ablation in animal model and cultured islets, could underlie the divergent outcomes between the two studies.

## Limitations of the study

While our study offered novel insights into sex-specific islet cell Ca²⁺ dynamics in response to glucose, key questions remain. First, ∼10% of insulin-positive cells expressed GCaMP6 in our mouse model. While these cells were identified as β-cells according to their Ca²⁺ dynamics, which resembled those of non-GCaMP-expressing β-cells within the same islets, we cannot entirely exclude the possibility that they represent a distinct α-cell subpopulation. Further work specifically characterizing this subpopulation may provide fundamental insights into α-/β-cell plasticity and potential transdifferentiation. Second, our *ex vivo* slice preparation is limited in capturing potential *in vivo* inputs such as innervation, blood flow, and circulating metabolites^61^. Thus, we can not exclude these factors as potential contributors to α-cell regulation. We also acknowledge that cytosolic Ca²⁺ changes may not fully represent hormone secretion in either β-^62^ or α-cells^63^, nevertheless, it remains a useful proxy for assessment of cellular function^35^. Third, since insulin and SST receptor antagonists were applied globally in intact pancreas slices, where native cell–cell interactions were preserved, blocking SST or insulin receptors may concurrently have altered input strength from neighboring β- or δ-cells, which expressed somatostatin and insulin receptors as well^41,64^. This complexity made it difficult to isolate direct effects of insulin or somatostatin on α-cells in pancreas slices. Nonetheless, our approach offered valuable mechanistic insights into islet α-, β-, and δ-cells interactions under both unperturbed and perturbed paracrine signaling in the most physiologically relevant *ex vivo* environments. Fourth, we cannot exclude inherent transcriptomic and proteomic heterogeneity among individual α-cells, including those showing β-cell-like Ca²⁺ responses to elevated glucose. While single-cell RNA-Seq^65^ supported functional heterogeneity in α-cells in pancreas slices, future studies using immunocytochemistry^66^ and *in situ* hybridization^67^ may offer an additional means to assess this possibility. Finally, it will be important in the future to extend our findings to human pancreas, where the higher proportion and intermingling between α-/β-cells and other islet cells^21^ may further emphasize paracrine regulation.

In summary, our study provided a simultaneous, cell type-specific characterization of Ca²⁺ dynamics in pancreatic α-, β-, and δ-cells in response to dynamic glucose changes. These findings revealed heterogeneity in α-cell Ca^2+^ dynamics, which is shaped both by glucose but also by insulin and glucagon secreted from neighboring β-, and δ-cells. We further identified sex- dependent differences in α-cell responses and heterogeneity, laying the groundwork for future investigations. Ultimately, these insights may contribute to the development of more targeted and effective strategies for diabetes treatment.

## Acknowledgements

We thank Dr. Aurelie Cleret-Buhot and the CRCHUM Cell Imaging, and the Animal and Cellular Physiology Facilities for their assistance.

## Author contributions

Y.-C.H. conceptualized the project, designed and conducted all experiments and performed data analyses of the study. Y.-C.H also created all figures and wrote the manuscript. G.A.R oversaw and supervised the research, supplied experimental resources and critically reviewed the manuscript. M.S.R. provided the analytical pipeline for imaging analysis, contributed critical consultation and mentorship on data analysis, script construction and critically reviewed the manuscript.

