## Supplementary Methods and Figure legend for "Pancreatic α-cells are functionally heterogeneous and sex-dependently regulated by neighboring endocrine cells"

*Blood glucose measurement, blood collection, IPGTT and ITT*

Blood glucose was measured from pre-nicked mouse tails using a glucometer (Ascensia, Mississauga, ON, Canada). For serum isolation, blood was collected from the pre-nicked tail using capillary tubes (KIMBLE DWK Life Science, USA), supplemented with HALT protease inhibitor (Thermo Fisher Scientific, Canada), and centrifuged at 10,000g for 10 minutes to collect the supernatant.  For IPGTT and ITT, mice were fasted either overnight for 16 hours or 4 hours, respectively, with ad libitum access to water. Baseline blood glucose levels were measured via tail vein sampling using a glucose meter. Glucose (2g/kg body weight) or Insulin (0.75 U/kg body weight, Novolin®ge Toronto, Novo Nordisk, ON, Canada) was diluted in sterile PBS and administered via intraperitoneal injection. Blood glucose levels were recorded at baseline (0 minute) and 15-, 30-, 60-, and 90-minutes post-injection.

*Pancreas slice preparation*

The pancreas slices (140 µm) were maintained in HEPES-ECS (in mM: 125 NaCl, 10 NaHCO3, 10 HEPES, 6 glucose, 6 lactic acid, 3 myo-inositol, 2.5 KCl, 2 Na-pyruvate, 2 CaCl2, 1.25 NaH2PO4, 1 MgCl2, 0.25 ascorbic acid; titrated to pH 7.4 using 1 M NaOH) or loaded with fluorescent $Ca^{2+}$ indicator in HEPES-ECS (Calbryte $590^{AM}$ (AAT Bioquest, CA, USA), 0.03% Pluronic F-127 (w/v), and 0.12% dimethylsulphoxide (v/v)).

*Immunofluorescence staining*

Once image acquisition was completed, pancreatic slices were immediately fixed in PBS containing 4% paraformaldehyde for 20 minutes, then permeabilized for 15 minutes at room temperature with 0.3% Triton X-100 in PBS. Islet containing pancreas slices were incubated overnight at 4 °C in PBS containing 0.3% Triton X-100, 5% bovine serum albumin, a primary monoclonal mouse anti-glucagon antibody (1:1000; Sigma-Aldrich, Canada), a polyclonal guinea pig anti-insulin antibody (1:10; DAKO, Santa Clara, CA, USA), and a monoclonal rat anti-somatostatin antibody (1:500; Invitrogen). The following day, pancreatic slices were PBS-washed and incubated for 4 hours at room temperature with secondary anti-rat, anti-guinea pig, and anti-mouse antibodies conjugated with Alexa Fluor 405, 561, and 647 (1:250 dilution). Slices were then PBS-washed and imaged immediately. GCaMP, Insulin (INS), Glucagon (GCG), Somatostatin (SST) and total islet area were determined using ZEISS ZEN 3.7 software on the acquired images.

*Pairwise Pearson correlation analysis*

Prior to analysis, traces were aligned in time and detrended to remove baseline fluctuations. The Pearson correlation coefficient measures the linear relationship between two time series according to the formula:

$$r = \frac{\sum_{i=1}^{n}(X_i - \bar{X})(Y_i - \bar{Y})}{\sqrt{\sum_{i=1}^{n}(X_i - \bar{X})^2}\sqrt{\sum_{i=1}^{n}(Y_i - \bar{Y})^2}}$$

where $X_i$ and $Y_i$ represent fluorescence values at time point $i$ from two cells, and $\bar{X}$ and $\bar{Y}$ are their respective mean fluorescence values. A coefficient of $r = 1$ indicates perfect positive correlation, $r = -1$ indicates perfect negative correlation, and $r = 0$ indicates no linear correlation.

*Network analysis*

We constructed functional α-cell networks from $Ca^{2+}$ dynamics using NetworkX in two steps. First, correlations c(i,j) for all pairs (i,j) of α-cells are obtained from recorded time series of $Ca^{2+}$oscillation. Second, these correlations were compared against a threshold ($c_0 = 0.7$): if $c(i,j)>c_0$, a line was placed between cells i and j. Networks were generated across six temporal segments corresponding to the glucose ramp. Connected nodes were visualized in red and isolated nodes in navy. For each network, we quantified mean degree (average number of links per node) and clustering coefficient (tendency of nodes to form local clusters). Mean degree values of 0–2 indicated sparse connectivity with mostly isolated cells; 3–7 indicated weakly connected networks with limited hubs; 8–20 reflected functional networks with coordinated communication; and >20 suggested highly synchronous networks. Clustering coefficients of 0–0.1 indicated disconnected clusters, 0.1–0.3 partial local coordination, 0.3–0.6 functional subnetworks, and 0.6–1.0 strong local hubs.

*Quantification of cell active fraction*

For each ROI, $Ca^{2+}$ events were grouped into consecutive 60-second intervals, and the total time during which $Ca^{2+}$ signals were elevated was calculated. The active fraction was defined as the total active time divided by 60 seconds for each interval. Median active fractions across each glucose concentration range for each cell type were then computed and plotted to compare the temporal activity dynamics of α- and δ-cell ROIs.

### Supplementary figure legends

**Supplementary Figure 1: Functional network analysis of α-cell $Ca^{2+}$ dynamics**

**A.** Representative α-cell networks in female (top) and male (bottom) islets across six glucose concentrations corresponding those in the glucose ramp. Each dot (known as nodes) represents an α-cell ROI. Connected nodes are shown in red, isolated nodes in navy. Note the weak connectivity across all networks with mostly isolated nodes. Quantification of **B.** Mean degree (average links per node) and **C.** clustering coefficient (local clustering tendency) across the six-glucose concentration over 30 islets from 5 females (85 ± 7.81 days) and 32 islets from 6 males (76 ± 9.31 days).

**Supplementary Figure 2: Spatial mapping reveals anticorrelated $Ca^{2+}$ activity between juxtaposed α- and β-cells in mouse islets**

**A.** A representative islet underwent high-speed calcium imaging followed by post-hoc immunoassay. The image shows GCaMP expression in glucagon-positive cells and their spatial relationship to insulin-positive areas. The corresponding glucagon- and insulin-positive regions are spatially mapped in **B.** Scatter plots display α-cells (blue) and β-cells (orange) positioned according to their x–y coordinates within image **A**. Negatively correlated α- and β-cell ROIs are spatially mapped. Lines connecting α–β cell pairs represent Pearson correlation coefficients (r) of $Ca^{2+}$ activity, color-coded by strength according to the scale bar on the right. **C.** This heatmap visualizes Pearson correlation coefficients computed between $Ca^{2+}$ activity traces of α-cell and β-cell ROIs within islet shown in **A.** The matrix is divided into four blocks by dashed lines: top-left for α–α correlations, bottom-right for β–β correlations, and the off-diagonal blocks for α–β and β–α correlations. Color intensity and hue represent correlation strength and direction, with red indicating positive and blue indicating negative correlations, centered at zero.

**Supplementary Figure 3: Quantification of active fraction of adjacent α- and δ-cells and heterogeneous $Ca^{2+}$ activity in GluCre:GCaMP6f mouse**

**A.** Each of the four panels shows the active fraction of individual cells, corresponding to the quantification of $Ca^{2+}$ activity trances of α- and δ-cell pairs in the main Figures 2E-1, 2F-2, 2F-1, and F-2 (also labeled at the top left of each panel). Green triangles (for δ-cells) and red circles (for α-cells) indicate individual measurements of active fraction per 60 seconds. Active fraction is defined as the proportion of each 60-second interval during which $Ca^{2+}$ signals were elevated. The overlayed dark green triangles and dark red circles highlight the median active fraction across each glucose concentration range. Cell types and ROI numbers are indicated in the legends. Note the inverse relationship in the $Ca^{2+}$ activity between α- and δ- cells in the left panels, and the temporally inverse relationship in the right panels.

**B.** Left panels show islets from four different mice, immunoassayed for glucagon (GCG) and somatostatin (SST) to identify α- and δ-cells, respectively. Right panels display the corresponding $Ca^{2+}$ activity traces recorded from δ- cells within the islets shown on the left. Note the heterogeneous nature of $Ca^{2+}$ dynamics among individual δ-cells and across different islets.

**Supplementary Figure 4: Somatostatin suppresses α-cell $Ca^{2+}$ dynamics via SSTR-mediated inhibition in female mice.**

Somatostatin (SST; 10, 50, 100nM, 10 minutes for each concentration), when applied to female **(A)** and male **(B)** GluCre:GCaMP6f mouse pancreas slices at 3.6mM glucose condition, reduced the frequency of $Ca^{2+}$ activities in α-cells in a dose dependent manner. Each circle (pink: female; blue: male) represents the mean α-cell $Ca^{2+}$ activity frequency of a single islet. Insets show quantification of mean ± SEM from individual islets represented in the main plots. Two-way ANOVA for repeated measures was performed. $^{*}p < 0.05$; $^{**}p < 0.01$. Note the steeper slope (orange dashed line) in **A-inset** than in **B-inset,** indicating greater somatostatin sensitivity in female α-cells. n = 9 islets from 3 female mice (101 ± 6.2 days) and 9 islets from 3 male mice (113.3 ± 1.5 days).

Somatostatin receptor antagonists MK4256 and CYN154806 (SSTRA; 400 and 300 nM, respectively, 10 minutes for each concentration) increased α-cell $Ca^{2+}$ activity in females (**C**; 10 islets from 3 mice, 101 ± 6.2 days), but only in few male islets (**D**; 9 islets from 3 mice, 113.3 ± 1.5 days). Two-way ANOVA for repeated measures was performed. $*p < 0.05$. In female islets, SSTRA-induced potentiation was reversed by 50 or 100nM SST, suggesting binding competition between SST and SSTRA and confirming a direct regulatory effect of somatostatin on α-cell excitability.

**Supplementary Figure 5: Effects of insulin and insulin receptor blockade on α-cell $Ca^{2+}$ dynamics.**

Insulin (INS; 200, 2,000, 20,000μU/mL) was applied to pancreas slices from female **(A)** and male **(B)** GluCre: GCaMP6f mouse at 3.6mM glucose condition, resulting in a downward trend in α-cell $Ca^{2+}$ activity frequency **(A and B, insets)**. The insulin receptor antagonist S961 (10, 100, 500) elicited a mildly potentiated of α-cell $Ca^{2+}$ dynamics in both sexes **(C, females; and D, males)**. Each data point (pink circle: female; blue circle: male) represents the mean α-cell $Ca^{2+}$ oscillation frequency for an individual islet. Insets display quantification (mean ± SEM) from islets shown in the main panels. INS treatment: 10 islets from 3 females (86 ± 36 days) and 11 islets from 3 males (93 ± 5.1 days). S961 treatment: 11 islets each from 3 female and 3 male mice (86 ± 36 and 93 ± 5.1 days, respectively). Two-way ANOVA for repeated measures was performed.

**Supplementary Figure 6: Population-level $Ca^{2+}$ activity dynamics visualized by hexbin event density plots.**

Hexbin density plots illustrate the spatiotemporal distribution of $Ca^{2+}$ events in α- and β-cells across all recorded islets. Panels **A-D** share a consistent layout: α-cells (left) and β-cells (right) from female (top row) and male (bottom row) GluCre:GCaMP6f mice were analyzed under the following conditions: (**A**) glucose ramp alone, (**B**) SSTR antagonists (SSTRA) + insulin receptor antagonist S961, (**C**) SSTRA alone, and (**D**) S961 alone. The x-axis indicates the time (in seconds) when $Ca^{2+}$ events reached their peak; the y-axis indicates event duration at half-maximum amplitude (halfwidth, log-scaled seconds). Warmer colors denote higher event density. These plots represent a global view of event timing and variability across the entire α- and β-cell populations.

**A)** Glucose ramp alone: α-cells: 30 islets from 5 female mice (85 ± 7.8 days) and 48 islets from 9 male mice (78.9 ± 10.6 days); β-cells: 21 islets from 5 female mice (85 ± 7.8 days) and 26 islets from 8 male mice (79.8 ± 8.8 days).

**B)** SSTRA + S961: α-cells: 19 islets from 3 female mice (88 ± 7.0 days) and 19 islets from 3 male mice (94.3 ± 6.4 days); β-cells: 16 islets from 3 female mice (88 ± 7.0 days) and 18 islets from 3 male mice (94.3 ± 6.4 days).

**C)** SSTRA alone: α-cells: 18 islets from 3 female mice (65.3 ± 5.5 days) and 18 islets from 3 male mice (77.3 ± 3.8 days); β-cells: 17 islets from 3 female mice (65.3 ± 5.5 days) and 11 islets from 3 male mice (77.3 ± 3.8 days).

**D)** S961 alone: α-cells: 19 islets from 3 female mice (84 ± 2.0 days) and 19 islets from 3 male mice (86.5 ± 10.1 days); β-cells: 16 islets from 3 female mice (84 ± 2.0 days) and 19 islets from 3 male mice (86.5 ± 10.1 days).
