## Supplementary Figures for "Pancreatic α-cells are functionally heterogeneous and sex-dependently regulated by neighboring endocrine cells"

### Supplementary Figure 1

A.

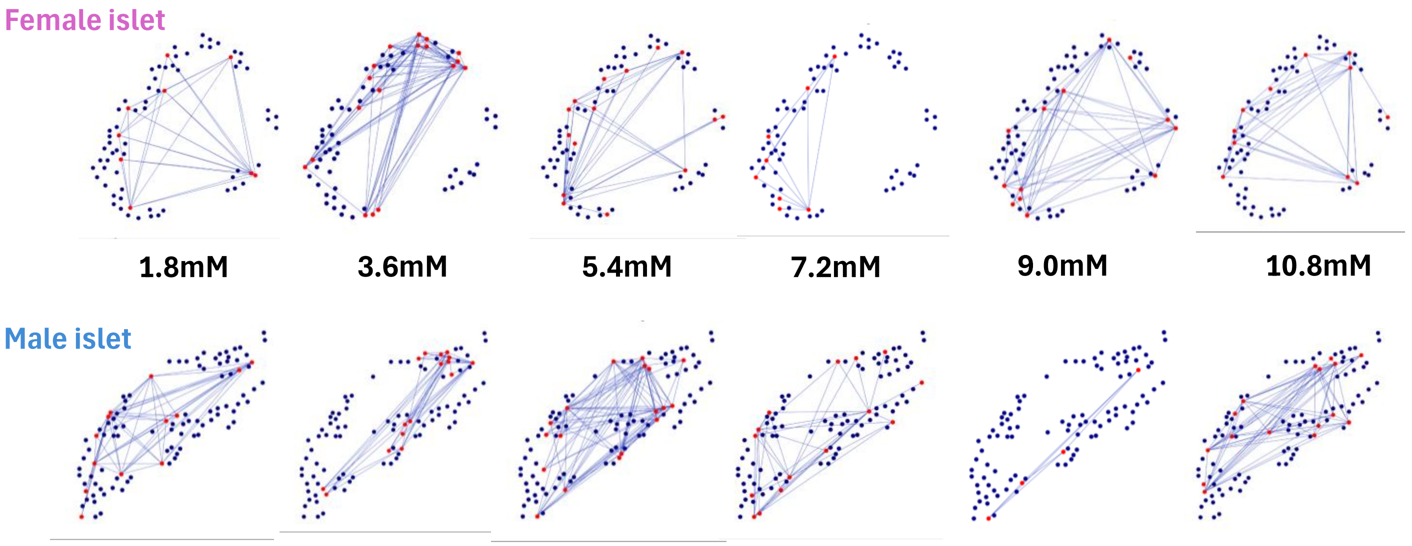


B.

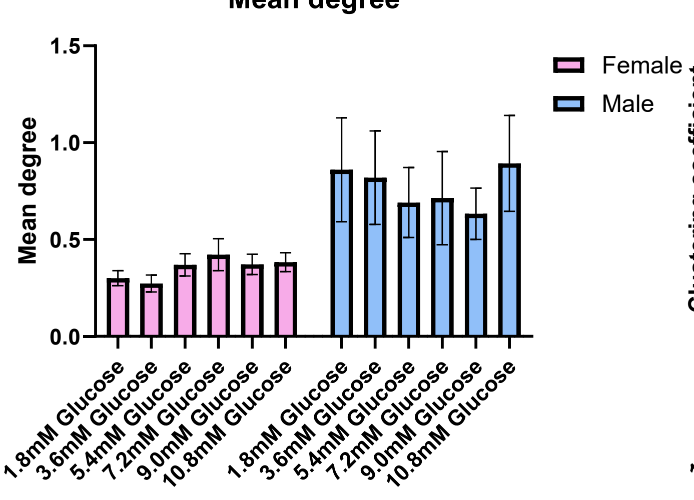


C.

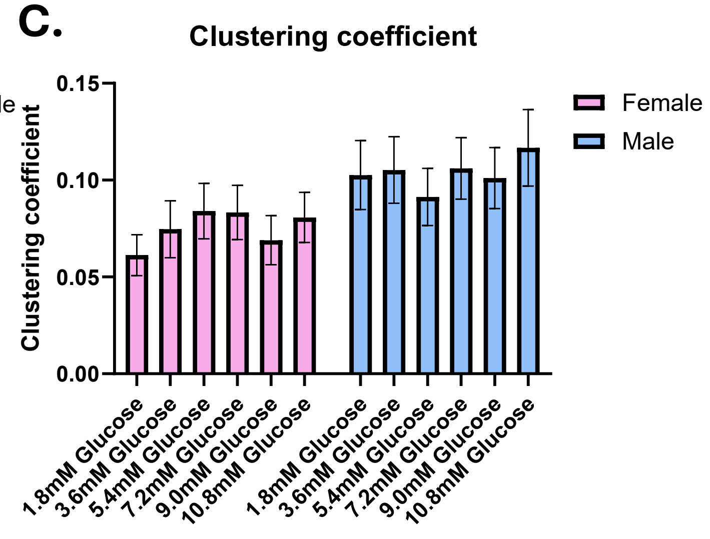

### Supplementary Figure 2

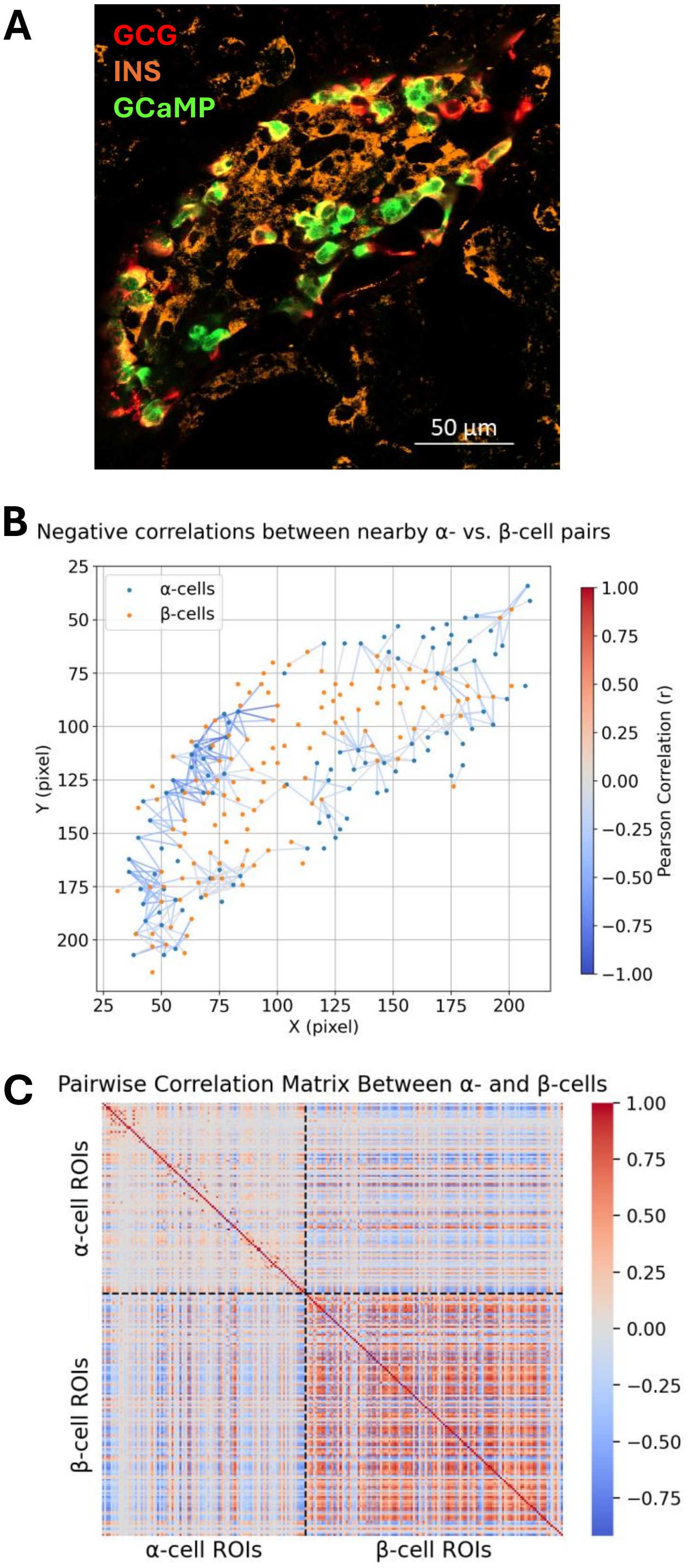

### Supplementary Figure 3

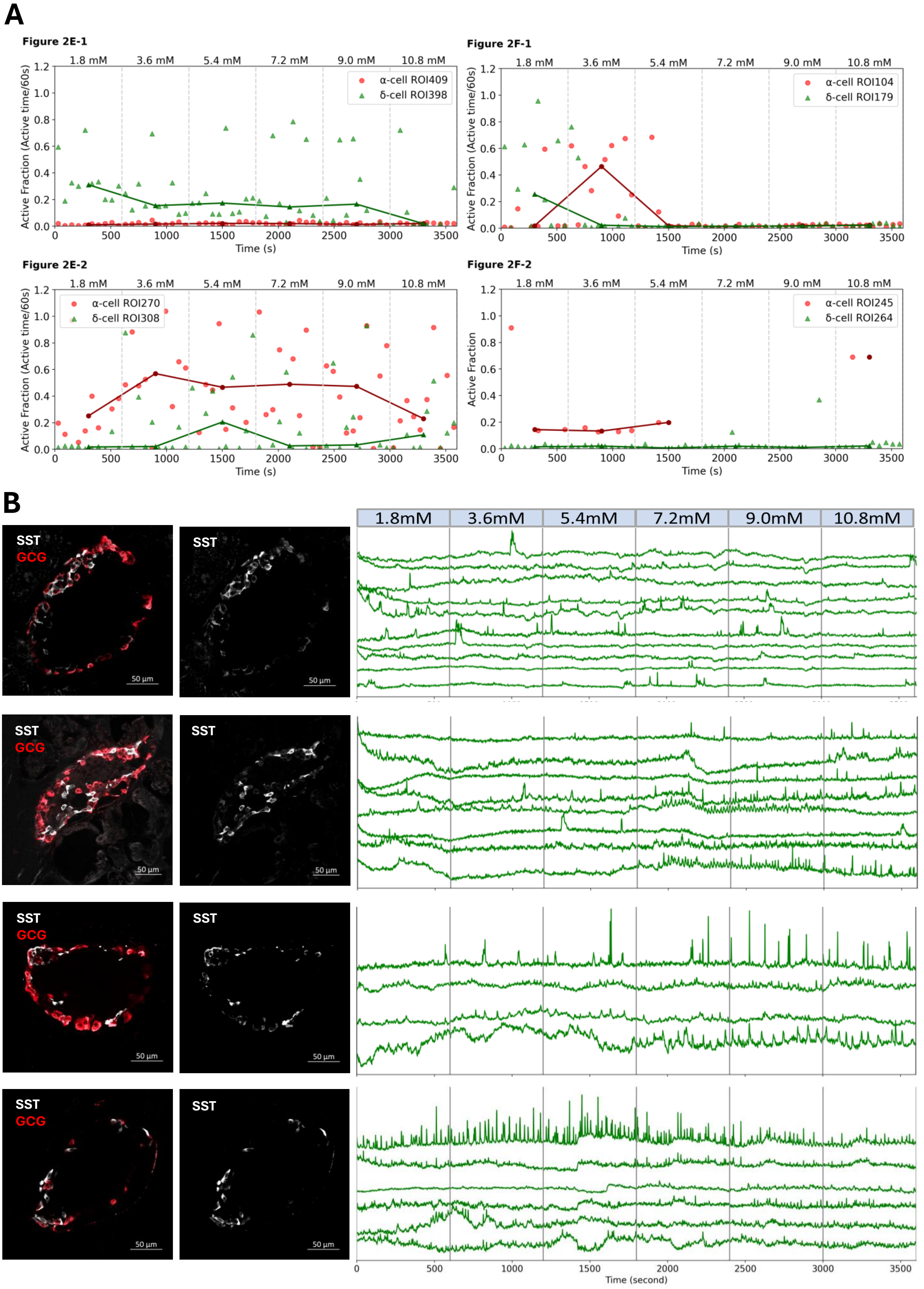

### Supplementary Figure 4

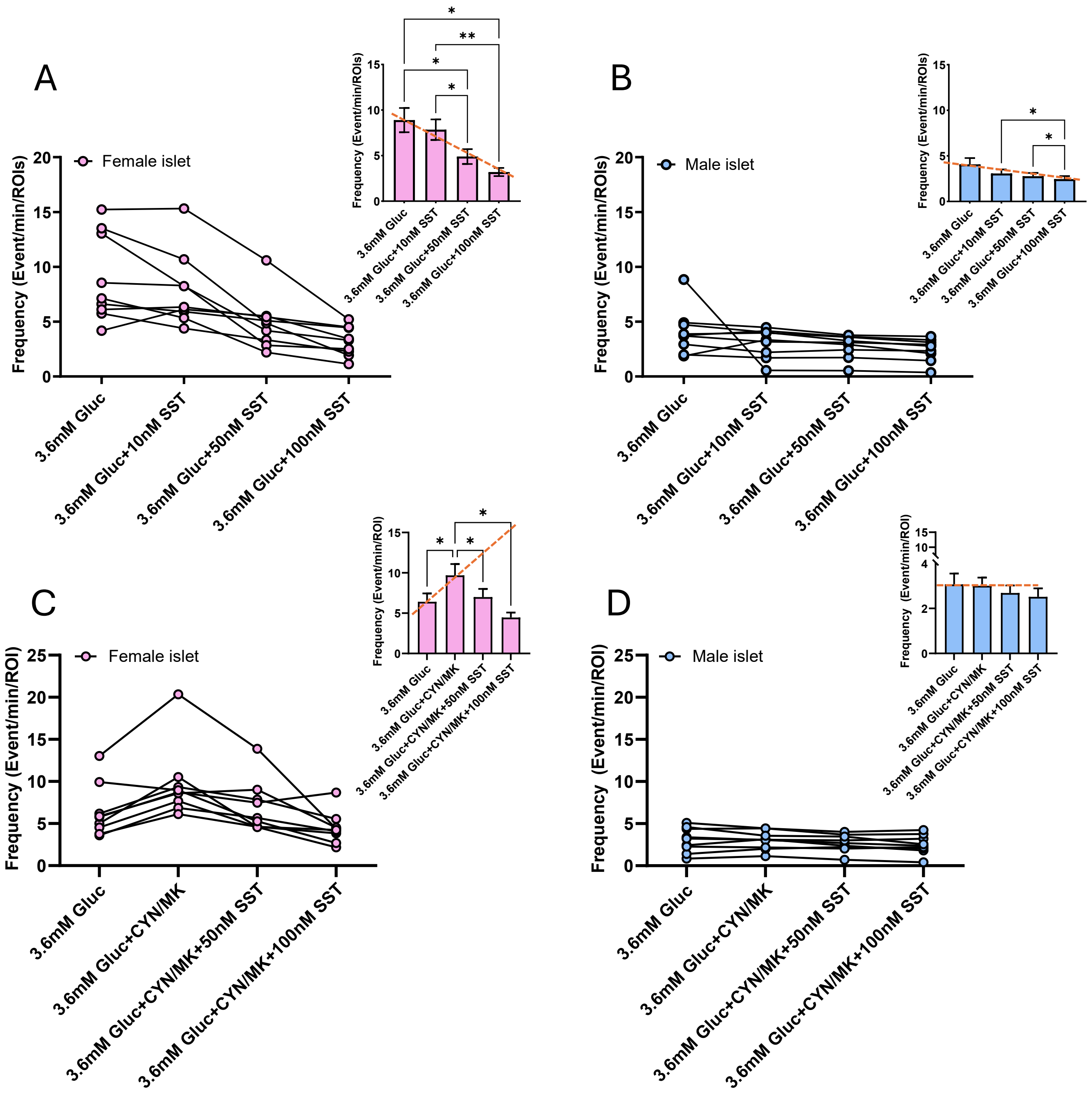

### Supplementary Figure 5

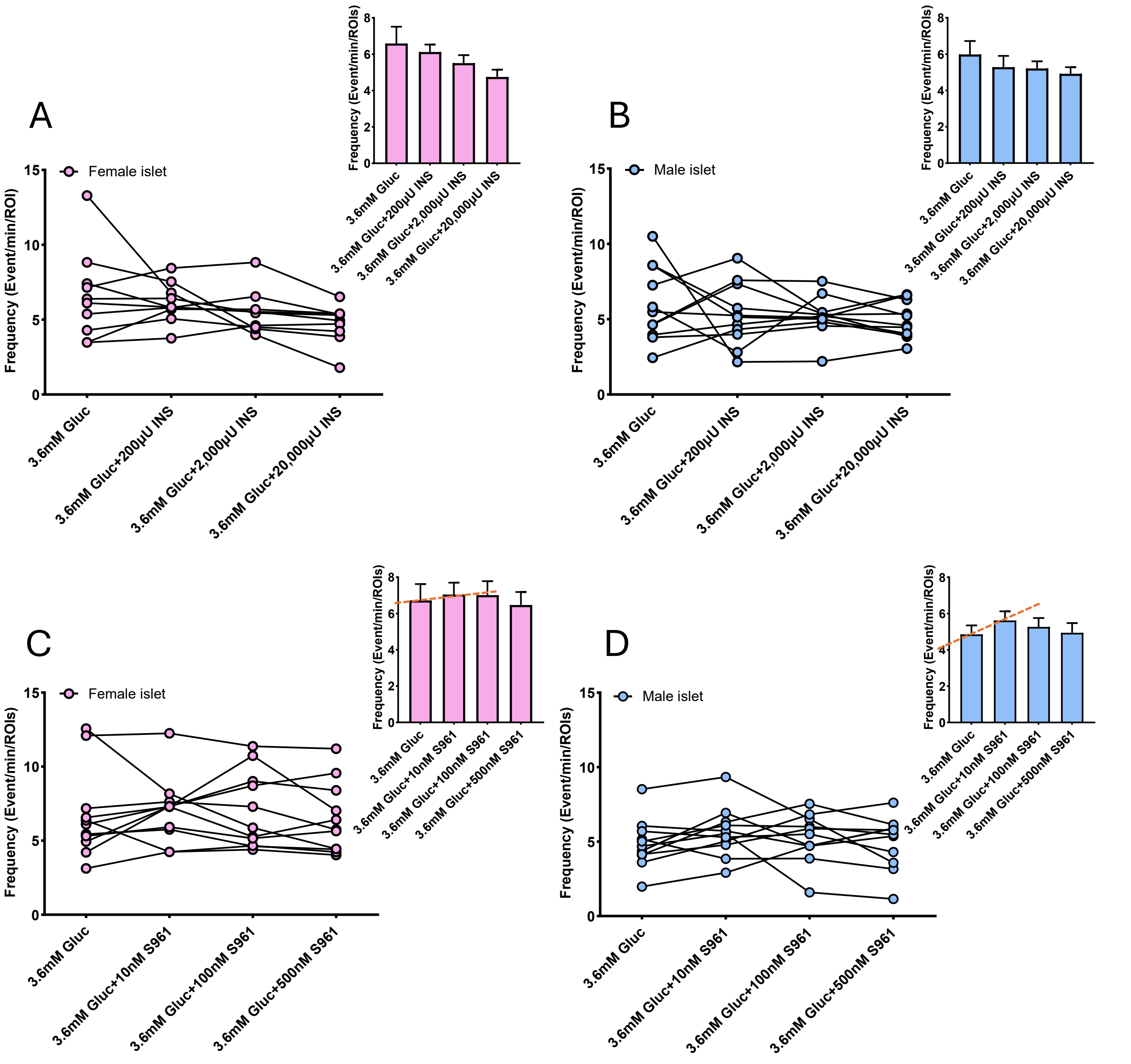

### Supplementary Figure 6

A)

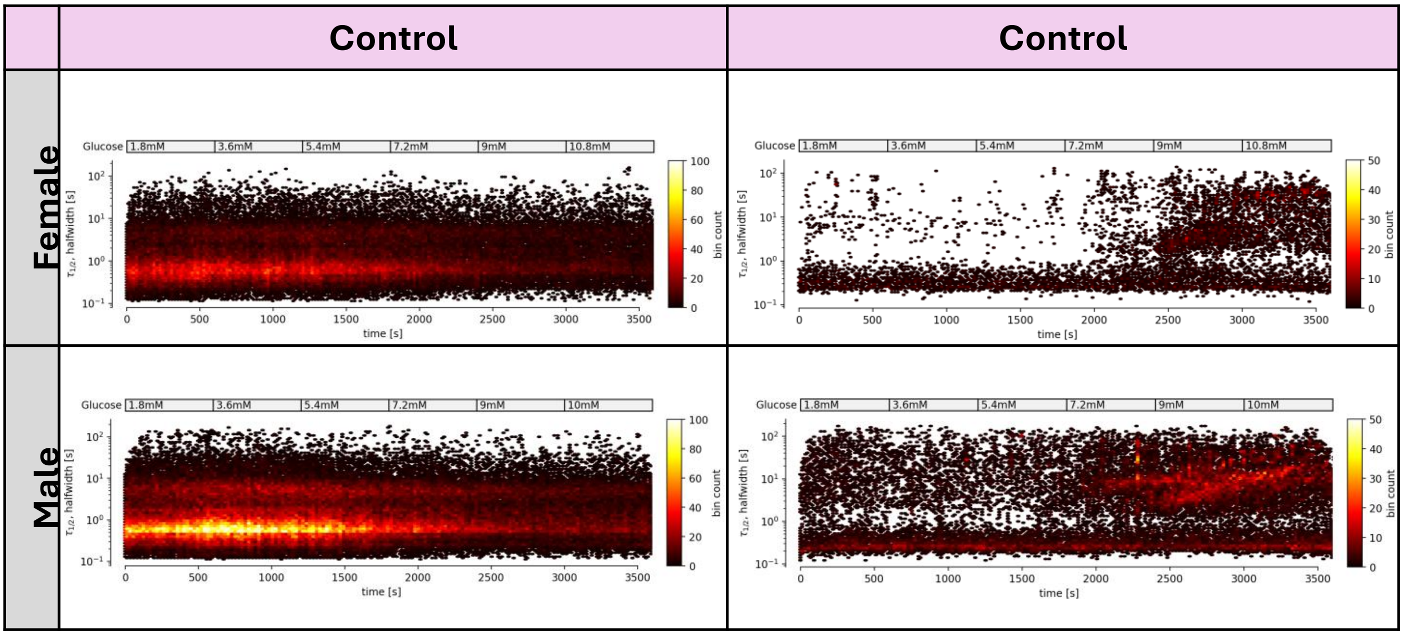


B)

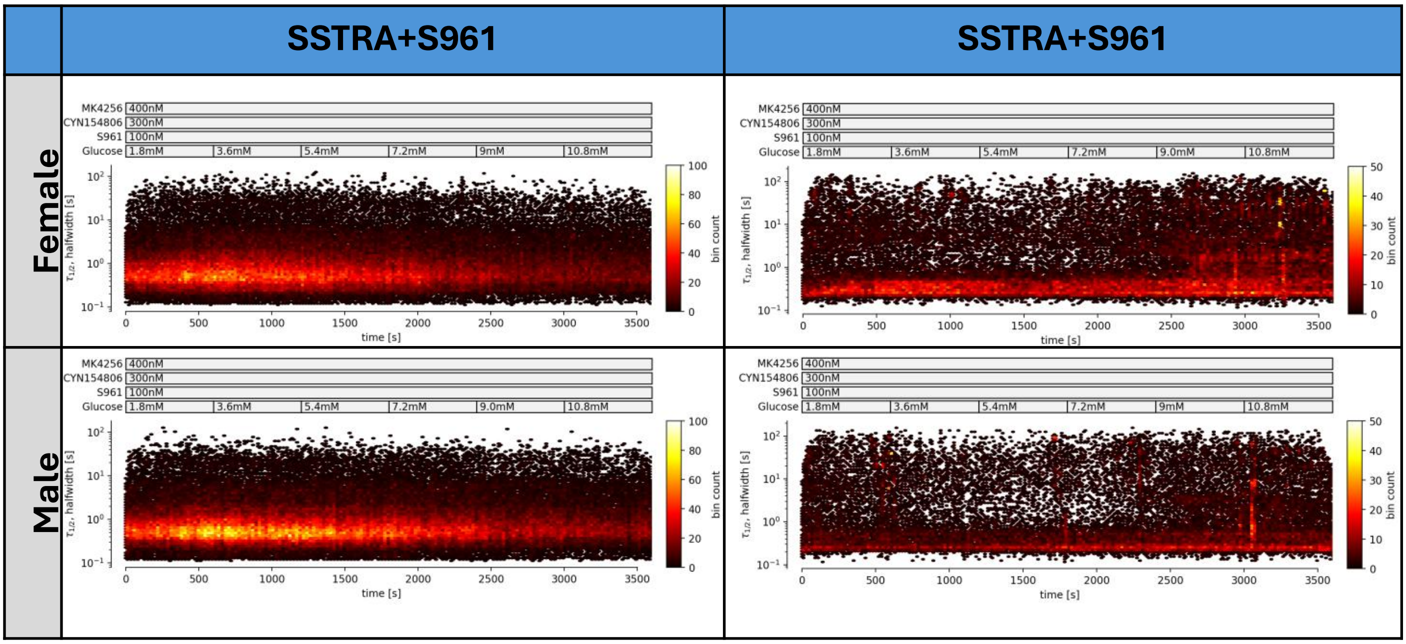

C)

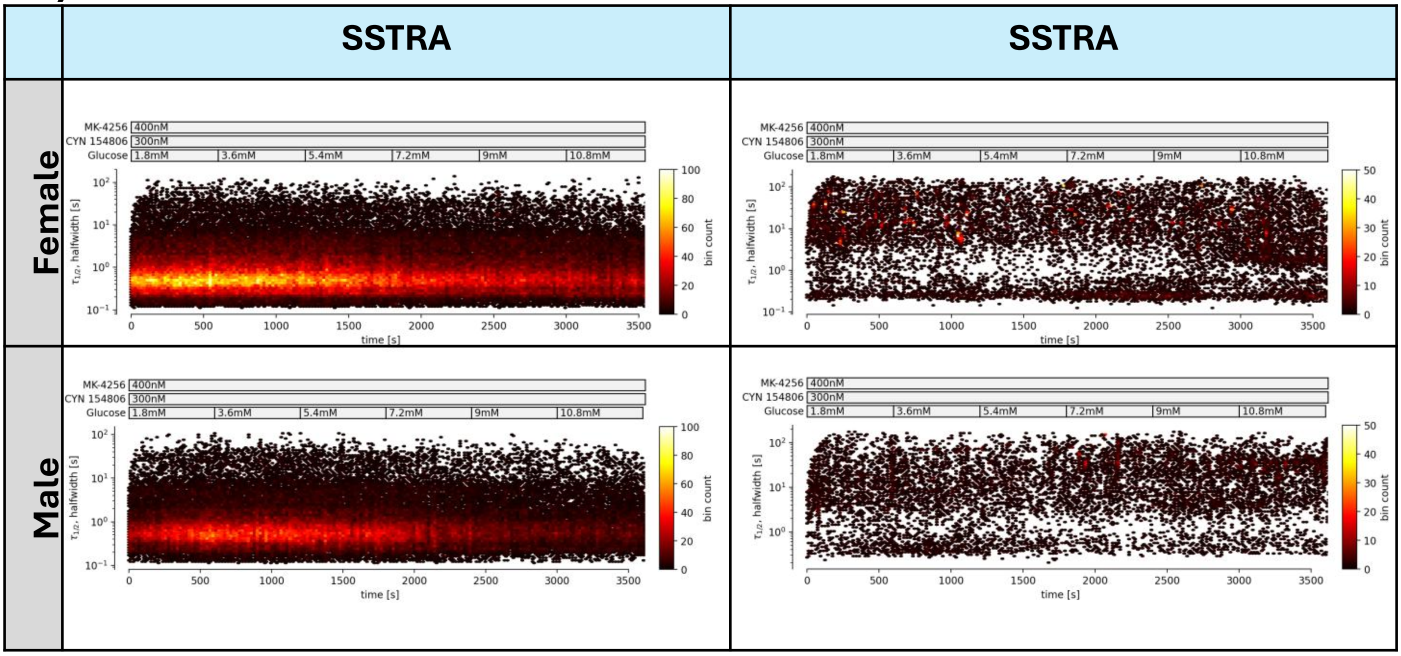

SSTRA
SSTRA
Female
Male
MK-4256 400nM
CYN 154806 300nM
Glucose 1.8mM 3.6mM 5.4mM 7.2mM 9mM 10.8mM
τ1/2, halfwidth [s]
time [s]
bin count


D)

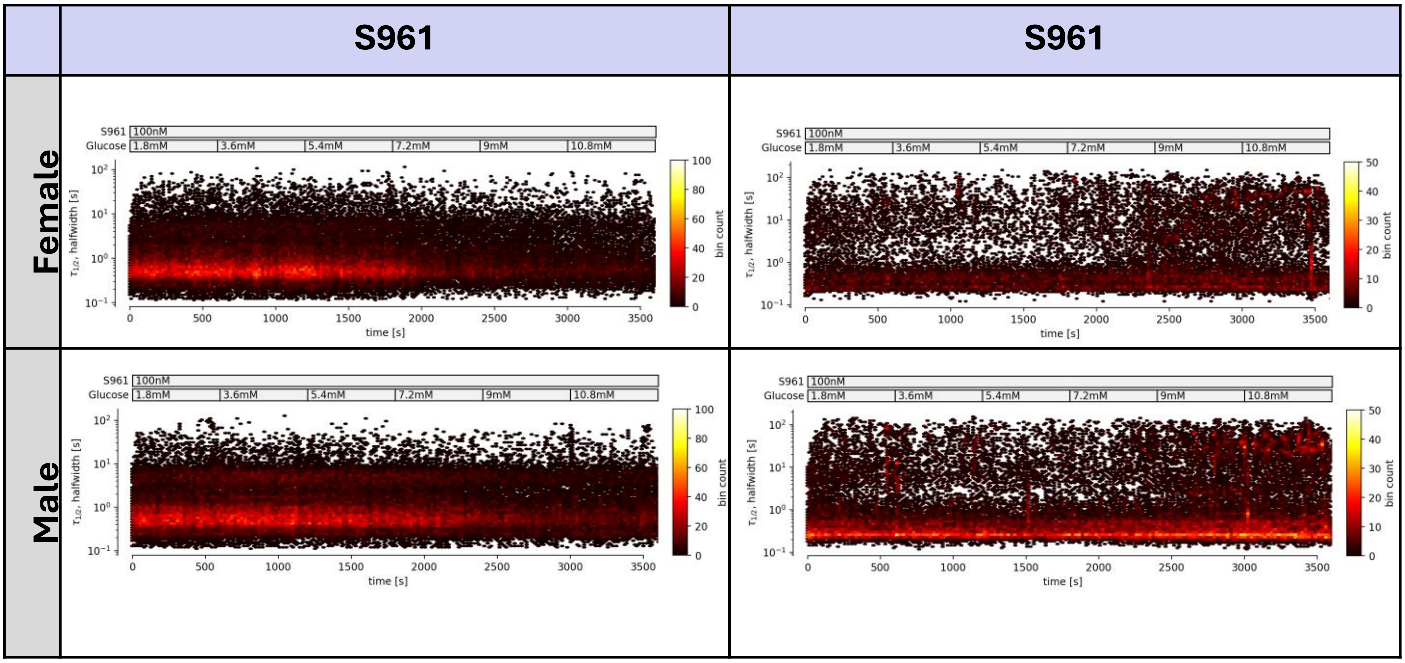

S961
S961
Female
Male
S961 100nM
Glucose 1.8mM 3.6mM 5.4mM 7.2mM 9mM 10.8mM
τ1/2, halfwidth [s]
time [s]
bin count
